# Glutamine Transport via Neurotransmitter Transporter 4 (NTT4, SLC6A17) Maintains Presynaptic Glutamate Supply at Excitatory Synapses in the Central Nervous System

**DOI:** 10.1101/2024.10.20.616835

**Authors:** Angela L. Nicoli, A. Shaam Al Abed, Sarah R. Hulme, Abhijit Das, Gregory Gauthier-Coles, Angelika Bröer, Sarojini Balkrishna, Gaetan Burgio, Nathalie Dehorter, Caroline D. Rae, Stefan Bröer, Brian Billups

## Abstract

The glutamate-glutamine cycle is thought to be the principle metabolic pathway that recycles glutamate at excitatory synapses. In this cycle, synaptically released glutamate is sequestered by astrocytes and converted to glutamine before being returned to the presynaptic terminal for conversion back into glutamate to replenish the neurotransmitter pool. While many aspects of this cycle have been extensively studied, a key component remains unknown: the nature of the transporter responsible for the presynaptic uptake of glutamine. We hypothesise that neurotransmitter transporter 4 (NTT4/*SLC6A17*) plays this role. Accordingly, we generated NTT4 knockout mice to assess its contribution to presynaptic glutamine transport and synaptic glutamate supply. Using biochemical tracing of [1-^13^C] glucose and [1,2-^13^C] acetate in awake mice, we observe a reduction of neuronal glutamate supply when NTT4 is absent. In addition, direct electrical recording of hippocampal mossy fibre boutons reveals a presynaptic glutamine transport current that is entirely inhibited by genetic or pharmacological elimination of NTT4. The role of NTT4 in neurotransmission was demonstrated by electrophysiological recordings in acute hippocampal slices, which reveal that NTT4 is required to maintain vesicular glutamate content and to sustain adequate levels of glutamate supply during periods of high-frequency neuronal activity. Finally, behavioural studies in mice demonstrate a deficit in trace fear conditioning; a hippocampus-dependent memory paradigm, and abnormalities in nest building, anxiety behaviour, and social preference. These results demonstrate that NTT4 is a presynaptic glutamine transporter which is a central component of the glutamate-glutamine cycle. NTT4 and hence the glutamate-glutamine cycle maintain neuronal glutamate supply for excitatory neurotransmission during high-frequency synaptic activity, and are key regulators of memory retention and normal behaviour.

## Introduction

In order to sustain ongoing synaptic transmission, presynaptic terminals must maintain an adequate neurotransmitter supply. At glutamatergic synapses, a significant portion of the released transmitter is thought to be recycled back to the presynaptic terminals indirectly, via astrocytes, by means of the glutamate-glutamine cycle (van den Berg and Garfinkel, 1971; Benjamin and Quastel, 1972; Berl and Clarke, 1983; Ottersen et al., 1992).

### The glutamate glutamine cycle

Following synaptic release, glutamate is ultimately removed from the extracellular space by high-affinity excitatory amino acid transporters (EAATs), which are known to be expressed predominantly on astrocytes rather than neurons (Rothstein et al., 1994; Chaudhry et al., 1995; Furness et al., 2008; Petr et al., 2015). A substantial proportion of this glutamate is then condensed with ammonia by the glial enzyme glutamine synthetase to form glutamine (Norenberg and Martinez-Hernandez, 1979; Rothstein and Tabakoff, 1984). Astrocytes release this glutamine back to the extracellular space by a variety of pathways (Deitmer et al., 2003), including a transporter mediated process involving e.g. Sodium-coupled Neutral Amino Acid Transporter 3 (SNAT3 / *Slc38a3*) (Todd et al., 2017; Dong et al., 2018) and a gap junction / hemichannel (connexin43) mediated release (Cheung et al., 2022). This extracellular glutamine is proposed to be transported into the presynaptic terminal, to be deamidated by phosphate activated glutaminase (Kvamme et al., 2001) for the resupply of presynaptic glutamate (Rae et al., 2003), thus forming the glutamate-glutamine cycle. Despite the elegance of this biochemical pathway for clearing synaptically released glutamate and resupplying the presynaptic neurotransmitter stores, its importance in maintaining excitatory neurotransmission is unclear. Up to approximately 50% of astrocytic glutamate is metabolised via the TCA cycle, rather than being returned to the presynaptic terminal (McKenna, 2007), and alternative presynaptic glutamate recycling / production pathways are potentially involved in replenishing neurotransmitter supply (Maciejewski and Rothman, 2008). Despite this, it is apparent that the glutamate-glutamine cycle plays at least a partial role in resupplying glutamate for excitatory neurotransmission (Rae et al., 2003; Tani et al., 2010; Billups et al., 2013; Tani et al., 2014).

### Presynaptic glutamine transport

The movement of glutamine into the presynaptic terminal is an essential component of the glutamate-glutamine cycle. However, the identity of the plasma membrane transporter responsible for this process is presently unknown. Therefore, to further characterise the glutamate-glutamine cycle and to investigate its role in maintaining levels of excitatory neurotransmission, we sought to identify and specifically inhibit this presynaptic glutamine transport process. Previous work using targeted recordings directly from the calyx of Held terminal has shown that presynaptic glutamine transport is electrogenic, relying on the transmembrane gradient of sodium to power glutamine transport into glutamatergic terminals. This transporter was also shown to reside on presynaptic vesicles and be inserted into the plasma membrane in a SNARE-dependent manner and retrieved by clathrin-mediated endocytosis, controlled by levels of synaptic activity (Billups et al., 2013). Since Neurotransmitter Transporter 4 (NTT4) is closely related to other presynaptic transporters (Bröer and Gether, 2012), is known to mediate the electrogenic, sodium-dependent transport of glutamine (Zaia and Reimer, 2009) and has been localised to neuronal presynaptic terminals, including localisation on a presynaptic vesicle pool (Masson et al., 1999), we hypothesised that NTT4 was the transporter responsible for presynaptic glutamine transport to support glutamatergic neurotransmission.

NTT4, also known as RXT1, is encoded by gene *SLC6A17* and is a member of the solute carrier 6 (SLC6) family of transporters, which includes GABA transporters and presynaptic transporters of other neurotransmitters such as noradrenaline, dopamine and serotonin (Bröer and Gether, 2012). Its substrates are mostly small neutral amino acids, including glutamine, proline, glycine, alanine and histidine, but notably also branched-chain amino acids (e.g. leucine) and the amino acid analogue α-(methylamino)isobutyric acid (MeAIB) (Parra et al., 2008; Zaia and Reimer, 2009). It is expressed exclusively in the central nervous system, with strong *in situ* hybridisation signals and immunohistochemical reactivity observed throughout the brain, including in the cortex, hippocampus, thalamus and cerebellum (Bröer, 2006). The physiological functions of NTT4 are presently uncertain. Sub-cellular localisation studies indicate that the majority of protein is located in presynaptic terminals on small vesicles (Fischer et al., 1999; Masson et al., 1999; Parra et al., 2008). Consistent with this observation is the suggestion that NTT4 is a vesicular transporter that loads glutamine into synaptic vesicles (Jia et al., 2023). However, here we present evidence that NTT4 functions as a plasma membrane transporter, which mediates glutamine influx into the presynaptic terminals to support glutamate production.

### Maintenance of glutamatergic neurotransmission

The general aim of this study is to understand the extent to which NTT4, and hence the glutamate-glutamine cycle more broadly, controls excitatory neurotransmission via the regulation of presynaptic glutamate supply. It is calculated that the presynaptic glutamate store is depleted within seconds to minutes of normal synaptic activity (Marx et al., 2015), therefore we hypothesise that removal or inhibition of NTT4 will result in an inability to sustain glutamatergic neurotransmission that is quickly apparent when synapses are activated. To investigate this, we have generated a new *Slc6a17^-/-^* mouse strain (NTT4-KO), which is deficient in glutamine transport. Using ^13^C tracing with NMR spectroscopy and liquid chromatography–mass spectrometry (LC-MS) we show decreased neuronal glutamate production in NTT4-KO mice. This is further supported by electrophysiological measurements of excitatory neurotransmission in mouse hippocampal brain slices, which establish that NTT4 is a plasma membrane glutamine transporter that is central to regulating the supply of glutamate for excitatory neurotransmission.

As sustained glutamatergic transmission is required for synaptic plasticity (Citri and Malenka, 2008) and hence normal memory processing and animal behaviour, we hypothesise that NTT4-KO mice will have deficits in these domains. Accordingly, we subjected mice to a range of behavioural tests and show that NTT4 and, by inference, a fully functioning glutamate-glutamine cycle is necessary for execution of regular behavioural tasks.

## Results

### Slc6a17 KO mice lack functional NTT4 protein

To investigate the role of NTT4 in regulating glutamate production and controlling animal behaviour, we created a new *Slc6a17*/NTT4 KO mouse strain. Western blots of forebrain synaptosomes, which contain presynaptic, postsynaptic and glial membranes, showed complete absence of the NTT4 protein in KO mice, whilst expression of B^0^AT2 (*Slc6a15*), EAAT1 (*Slc1a3*), and the actin and synaptophysin controls were unchanged (Figure 1A). In addition, uptake of the NTT4 substrates glutamine, leucine and proline were significantly reduced in synaptosomes from NTT4 KO animals compared to WT (Figure 1B). Synaptosomal glutamine uptake was reduced by 20.8 % (WT = 0.89 ± 0.03 of glutamate uptake, n = 4 vs KO = 0.70 ± 0.02 of glutamate uptake, n = 4, p < 0.01), leucine uptake reduced by 32.2 % (WT = 0.38 ± 0.02 of glutamate uptake, n = 4 vs KO = 0.26 ± 0.01 of glutamate uptake, n = 7, p < 0.001) and proline uptake reduced by 43.0 % (WT = 0.62 ± 0.05 of glutamate uptake, n = 4 vs KO = 0.35 ± 0.02, n = 8, p < 0.001). Immunofluorescence imaging showed NTT4 protein was highly expressed in *stratum lucidum/radiatum* in the hippocampal CA3 subfield of WT animals (Figure 1D) and was present to a lesser extent in both the *stratum pyramidale* and *stratum radiatum* in CA1 (Figure 1C). In contrast, NTT4 expression was absent in the KO animals (Figure 1E & F). Together, this demonstrates that NTT4 protein is present at synapses in WT animals, and is effectively deleted in NTT4 KO animals.

**Figure 1.**
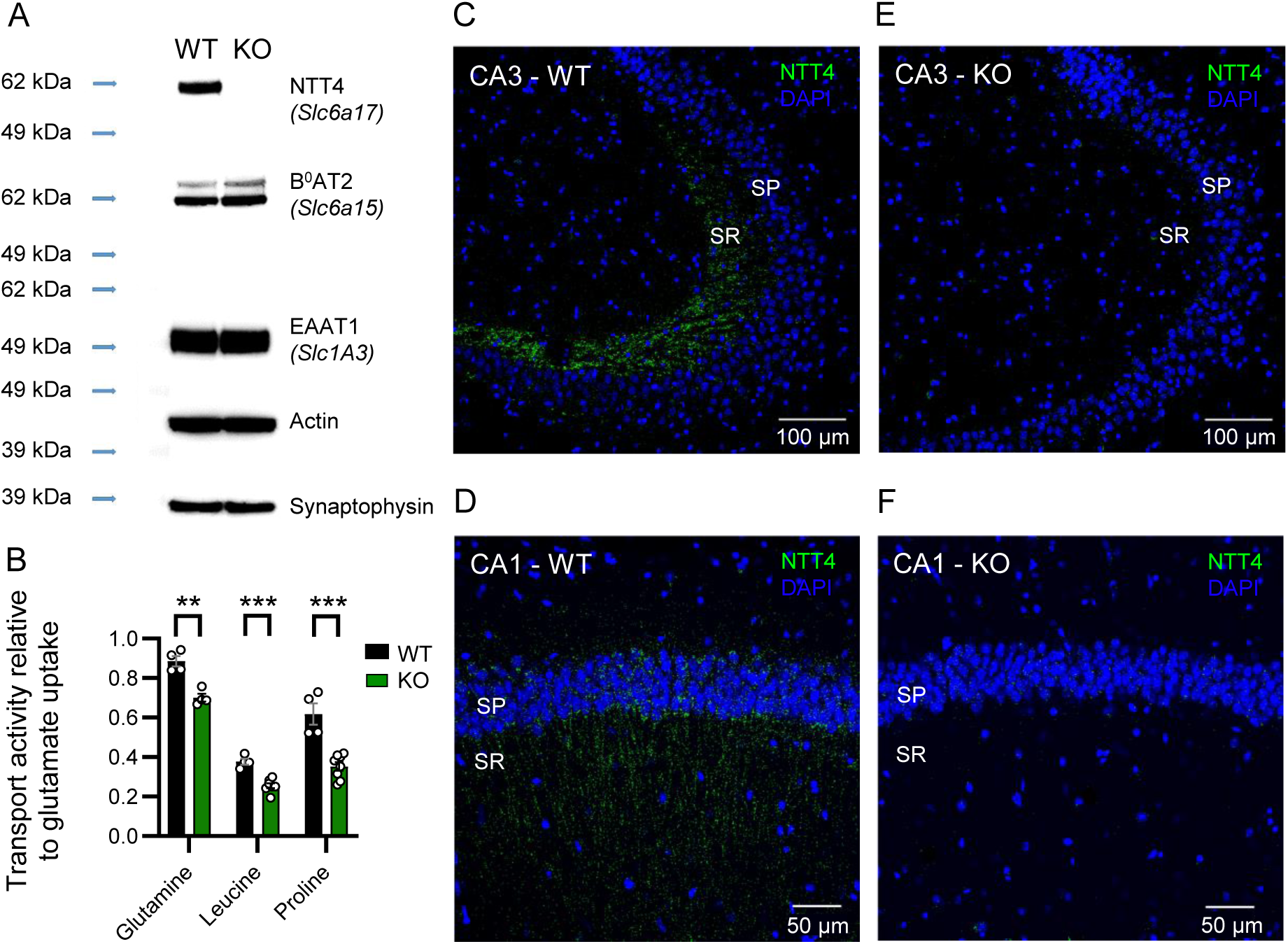
NTT4 knockout mouse (NTT4 KO; *Slc6a17^-/-^*) lacks functional NTT4 protein and shows reduced uptake of NTT4-substrate amino acids. **(A)** Western blot of synaptosomes prepared from forebrain tissue of WT and NTT4 KO mice shows NTT4 is absent NTT4 KO animals. There is no apparent upregulation of B^0^AT2 (*Slc6a15*), a transporter analogous to NTT4 with a similar substrate profile, or EAAT1 (*Slc1A3*), included here as a transporter control in the KO samples and both actin (loading) and synaptophysin (neuron) controls are present and unchanged. **(B)** Uptake of glutamine, leucine and proline relative to glutamate uptake in synaptosomes from WT (black) and NTT4 KO (green) mice. Synaptosomes from NTT4 KO animals have significantly reduced uptake of all three amino acids when compared to WT. ‘**’ indicates p < 0.01, ‘***’ indicates p < 0.001. **(C)** Immunohistochemistry in CA3 of WT mice shows NTT4 expression (green) is strong in *stratum lucidum/radiatum* (SR). **(D)** CA1 region in WT mice shows expression of NTT4 protein (green) in str*atum pyramidale* (SP), indicated by the presence of cell somas (DAPI = blue) and *stratum radiatum* (SR). **(E)** In CA3 of NTT4 KO animals, NTT4 expression (green) is absent. **(F)** In CA1 of NTT4 KO animals, NTT4 expression is absent.

### NTT4 KO mice have interrupted glutamate-glutamine flux

Flux of glutamate and glutamine in neurons and glia of awake, active mice was assessed by tracing of ^13^C enriched metabolic substrates. Following injection of WT and NTT4 KO mice with both [1-^13^C] glucose and [1,2-^13^C] acetate, mice were sacrificed, and labelled metabolites in cortex and hippocampus were quantified using ^13^C NMR (Figure 2A) and LC-MS (Figure 2B) respectively. Glutamate and glutamine derived from [1-^13^C] glucose (metabolised in both neurons and astrocytes) thus contain the single ^13^C at the C4 position whereas those produced via metabolism of [1,2-^13^C] acetate (metabolised in astrocytes (Rowlands et al., 2017)) are labelled at both C4 and C5.

**Figure 2.**
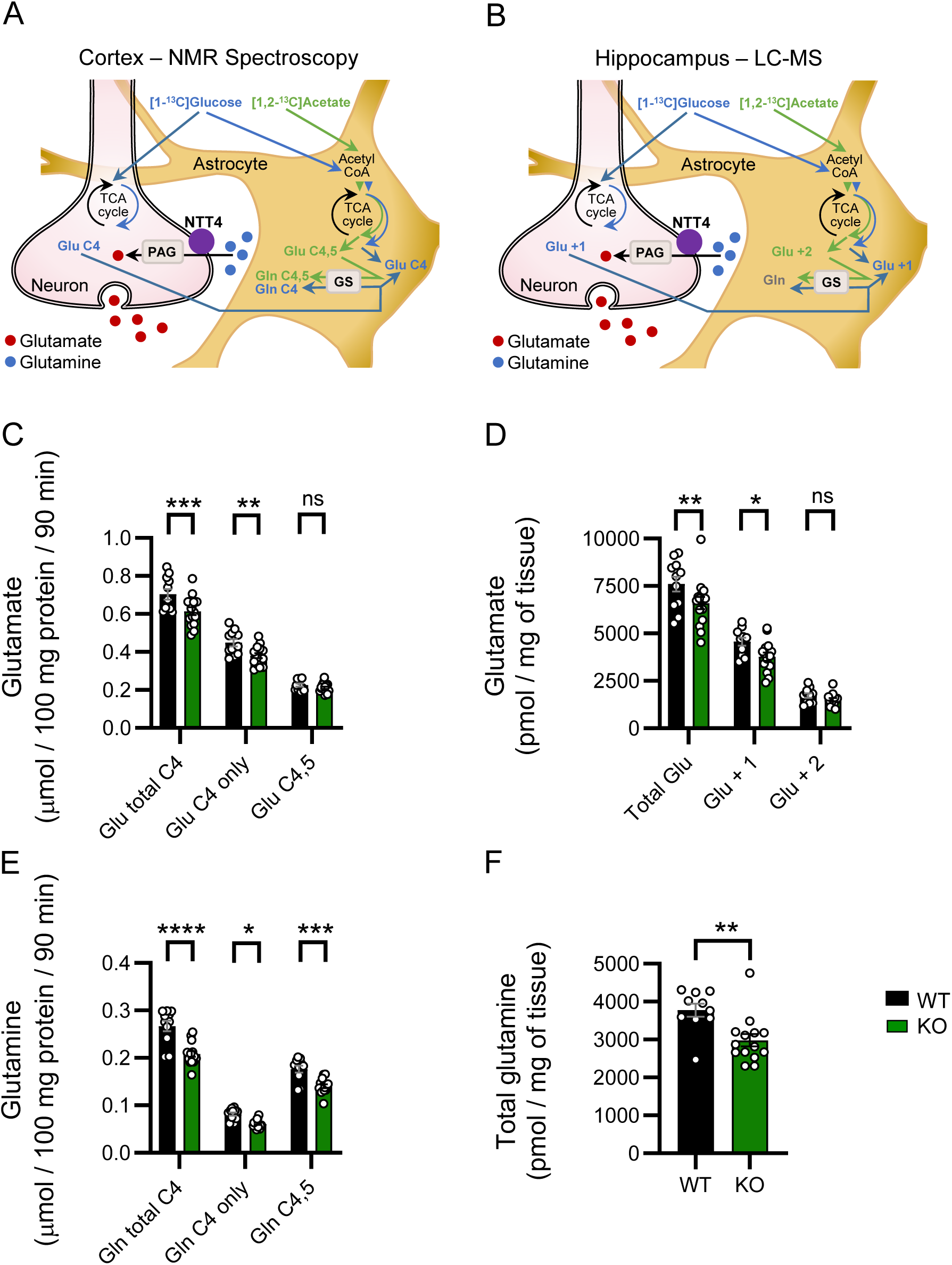
Glutamate and glutamine production are altered in NTT4 KO (*Slc6a17^-/-^*) mouse cortex and hippocampus. **(A)** Using ^13^C NMR spectroscopy, metabolism of [1-^13^C] and [1,2-^13^C] acetate into glutamate and glutamine was quantified in cortex of WT and NTT4 KO mice. Schematic diagram shows metabolism of isotopically enriched glucose via glycolysis in astrocytes and neurons (blue). Glucose-derived glutamate and glutamine are labelled only at C4. Isotopically enriched acetate (green) instead enters the TCA cycle via acetyl-coenzyme A (Acetyl-CoA) in astrocytes and is metabolised into glutamate and glutamine labelled at both C4 and C5 (C4,5). **(B)** Using LC-MS, metabolic products in hippocampus of WT (black) and NTT4 KO (green) mice were also quantified. Schematic diagram shows metabolism of isotopically enriched glucose and acetate as in A. For LC-MS quantification, the number of labelled carbons can be detected in glutamate, whereas glutamine quantification includes total labelled product. **(C)** Glutamate labelled only at C4 by metabolism of glucose is reduced in NTT4 KO mouse cortex, whereas acetate-derived astrocytic glutamate labelled at C4,5 is not. Total glutamate labelled at C4 is also reduced overall in NTT4 KO mice. N = 11 for WT and n = 15 for KO. **(D)** Similar to C, hippocampal glutamate derived from glucose (Glu +1) is reduced, whereas astrocytic glutamate labelled at two carbons (Glu + 2) is not. N = 10 for WT and n = 14 for KO. **(E)** In cortex, glutamine labelled only at C4 via metabolism of glucose is reduced in NTT4 KO animals, and so is glutamine labelled at C4,5 via astrocytic acetate metabolism. Accordingly, **(F)** shows total labelled glutamine is also reduced in hippocampus of NTT4 KO mice. ‘ns’ = no significant difference, ‘*’ = p < 0.05, ‘**’ = p < 0.01, ‘***’ = p < 0.001, ‘****’ = p < 0.000. Data presented at means ± SEM. Data points in C-F indicate individual mice.

Cortical samples analysed with ^13^C-NMR spectroscopy showed NTT4 KO mice had a reduced concentration of glutamate labelled only at C4 (WT = 0.45 ± 0.02 µmol / 100 mg protein / 90 min, n = 11 vs KO = 0.38 ± 0.01 µmol / 100 mg protein / 90 min, n = 15, p <0.01), but not at C4,5 (WT = 0.23 ± 0.01 µmol / 100 mg protein/ 90 min, n = 11 vs KO = 0.21 ± 0.01, µmol / 100 mg protein / 90 min, n = 15, p = 0.56) suggesting that the metabolism of glucose in either neurons or astrocytes, or both, was interrupted when NTT4 was absent. Conversely, the astrocytic metabolism of acetate was not affected. The reduction in C4-only labelled glutamate was significant enough to reduce the total glutamate with a label at C4, comprising both glucose and acetate-derived products combined (WT = 0.70 ± 0.03 µmol / 100 mg protein / 90 min, n= 11 vs KO = 0.61 ± 0.02 µmol / 100 mg protein / 90 min, n = 15, p < 0.001, Figure 2C).

Hippocampal samples analysed with LC-MS reflected a similar interruption, with glutamate containing a single ^13^C reduced in NTT4 KO mice (WT = 4568 ± 207 pmol / mg, n = 11 vs KO = 3775 ± 211 pmol / mg, n = 15, p < 0.05), whereas that containing two labelled carbons was no different (WT = 1722 ± 119 pmol / mg, n = 11 vs KO = 1510 ± 81 pmol / mg, n = 15, p = 0.55). Total ^13^C-labelled glutamate was also significantly different (WT = 7600 ± 392 pmol / mg, n = 11 vs KO = 6599 ± 323 pmol / mg, n = 15, p < 0.01, Figure 2D). Conversely, labelled GABA concentrations detected with ^13^C-NMR spectroscopy (Figure S1A) or LC-MS (Figure S1B) did not show any difference in NTT4-KO mice, showing that the reduction in glutamate levels is unlikely to have subsequent influence on inhibitory neurotransmission.

For glutamine, cortical samples from NTT4 KO mice had significantly lower incorporation of ^13^C at C4 only (WT = 0.08 ± 0.004 µmol / 100 mg protein / 90 min, n = 11 vs KO = 0.06 ± 0.002 µmol / 100 mg protein / 90 min, n = 15, p < 0.05) and at C4,5 (WT = 0.18 ± 0.01 µmol / 100 mg protein / 90 min, n = 11 vs KO = 0.14 ± 0.004 µmol / 100 mg protein / 90 min, n = 15, p < 0.001). This suggests that production of glutamine occurring in astrocytes both via metabolism of acetate, and by amidation of C4-labelled glutamate (derived from glucose), was impaired by the absence of NTT4. Accordingly, total C4 labelled glutamine was also significantly reduced (WT = 0.27 ± 0.01 µmol / 100 mg protein / 90 min, n = 11 vs KO = 0.21 ± 0.01 µmol / 100 mg protein / 90 min, n = 15 vs p < 0.0001, Figure 2E).

Glutamine analysed with LC-MS cannot be fragmented in the same way as glutamate, therefore the number of labelled carbons cannot be discerned. Consistent with the NMR results in cortical tissue however, hippocampal samples from NTT4 KO mice had less total labelled glutamine than control (WT = 3776 ± 173 pmol / mg, n = 10 vs KO = 2983 ± 166 pmol / mg, n = 14 vs, p < 0.01, Figure 2F).

### NTT4 mediates glutamine transport into presynaptic mossy fibre boutons

Mossy fibre boutons are the synaptic terminals of hippocampal dentate gyrus granule cells, which synapse onto CA3 pyramidal cells, and are seen to heavily express NTT4 (Figure 1D). As NTT4 is a sodium-coupled electrogenic transporter, it is possible to record transport currents when substrates are applied to the outside of voltage-clamped neurons if they express NTT4 at the cell surface. Thus, to evaluate the hypothesis that NTT4 is functional in the presynaptic terminal plasma membrane, we measured glutamine-induced transport currents in mossy fibre boutons in brain slices from WT and KO animals by direct patch-clamp recordings from the presynaptic terminal. Whole-terminal recordings at these boutons in slices from WT control animals revealed immediate inward currents induced by puff application of 10 mM glutamine (-2.4 ± 0.2 pA, n = 14, Figure 3A & C). To ensure that this glutamine-induced current is generated locally in the presynaptic terminal and is not the remnant of a larger current generated at the distal dentate gyrus granule cell soma, we also recorded glutamine transport currents directly from the granule cell soma. 10mM glutamine applied to the soma induced a current of -5.2 ± 0.6 pA (n = 8), which is not large enough to be the source of the presynaptic current we observe approximately 1 mm away. In contrast to WT slices, the presynaptic glutamine-induced current was absent in slices from NTT4 KO animals (-0.03 ± 0.08 pA, n = 6, p < 0.0001, Figure 3A & C). These data show that NTT4 sequesters glutamine locally at presynaptic terminals, and that in NTT4 KO animals, no glutamine transport currents remain in the presynaptic terminal membrane.

**Figure 3.**
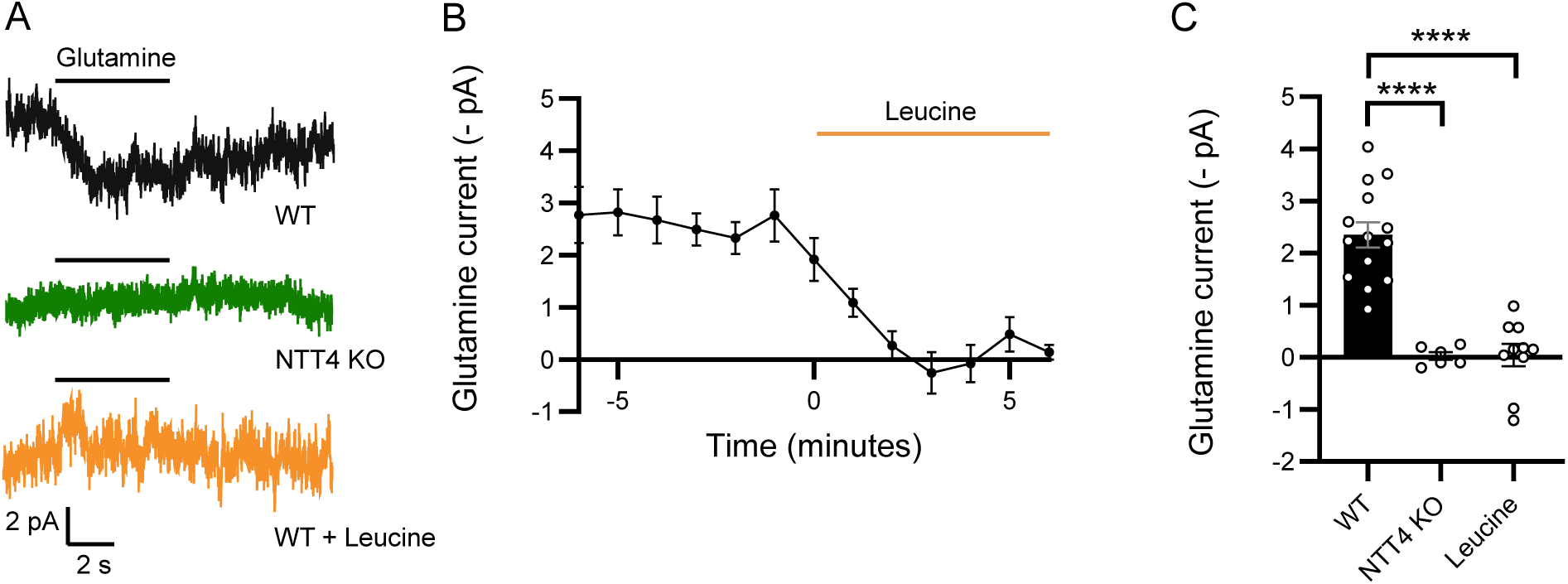
Presynaptic glutamine currents are abolished at mossy fibre boutons of NTT4 KO mice and when NTT4 is occluded. **(A)** Puff application of glutamine (‘Gln’) elicited inward currents in mossy fibre boutons in brain slices from WT animals (black trace, top). In NTT4 KO mice (green trace, middle) this current was no longer evoked by the presence of glutamine. Similarly, no current was elicited by the glutamine puff in WT brain slices when 10 mM leucine was present (orange trace, bottom). (B) Bath application of leucine abolished glutamine puff currents in mossy fibre boutons. The presence of leucine is indicated by the orange line. Data points show the mean peak amplitude of glutamine-evoked currents resulting from puff application of glutamine (5 s puff, repeated every minute) and error bars indicate SEM. (C) Mean puff-evoked glutamine current amplitude at mossy fibre boutons in WT, NTT4 KO and leucine-treated WT slices show currents present in WT control are absent in NTT4 KO and leucine-treated WT slices. Data points indicate individual recordings and error bars show SEM. ‘****’ indicates p < 0.0001.

There are currently no specific pharmacological inhibitors of NTT4, however the transporter has a higher affinity for leucine than for glutamine. Thus, application of a saturating concentration leucine will competitively block glutamine uptake. To see whether the glutamine uptake via NTT4 could be inhibited, leucine was applied to presynaptic terminals and responses to puff-applied glutamine measured. Following the addition of 10 mM leucine to the recording chamber, glutamine transport currents were quickly abolished in WT slices (reduced by 94 %, from -2.4 ± 0.3 pA to -0.05 ± 0.2 pA, n = 10, p < 0.001 Figure 3A-C). This validates the use of leucine as an acute substrate inhibitor of presynaptic glutamine transport. The glutamine induced current in the granule cell soma was unaffected by the addition of leucine (-4.9 ± 0.5 pA, n = 8, 96% of control, p = 0.29), indicating that NTT4 mediated glutamine transport is confined to the presynaptic compartment and glutamine transport in granule cell soma occurs via an alternative transporter, such as SNAT1 (*SLC38A1*) (Chaudhry et al., 2002b; Melone et al., 2004).

### NTT4 is required for rapid ongoing synaptic transmission at mossy fibre - CA3 synapses

Having demonstrated that NTT4 provides glutamine to the presynaptic terminal, we hypothesised that this glutamine acts as a precursor for synaptically released glutamate, and that inhibition of transport will hence reduce levels of glutamatergic neurotransmission. To measure glutamate release from synaptic terminals we firstly recorded EPSCs from CA3 pyramidal cells, while electrically stimulating mossy fibre inputs. To avoid recurrent excitation and kainate receptor mediated plasticity, the AMPA/kainate receptor antagonist NBQX (20 µM) was added to the bath, and glutamate release was assessed by recording NMDA receptor mediated currents. To remove the voltage-dependent Mg^2+^ block of NMDARs, cells were voltage-clamped at +20 mV, resulting in an outward NMDAR mediated EPSC, which was used as a measure of glutamate release. The role of NTT4 in replenishing glutamate during a low level of baseline synaptic activity was assessed by applying low-frequency stimulation (LFS; paired pulses 40 ms apart, repeated every 20 s, Figure 4A- top) and adding 10 mM leucine to the circulating ACSF. Perhaps surprisingly, EPSC amplitudes were unaffected by the addition of leucine for the duration of recording (from 39.0 ± 7.3 pA to 37.4 ± 7.3 pA n = 4, p = 0.13, Figure 4B & C) indicating that baseline synaptic activity can continue for an extended period in the absence of NTT4-mediated glutamine transport.

**Figure 4.**
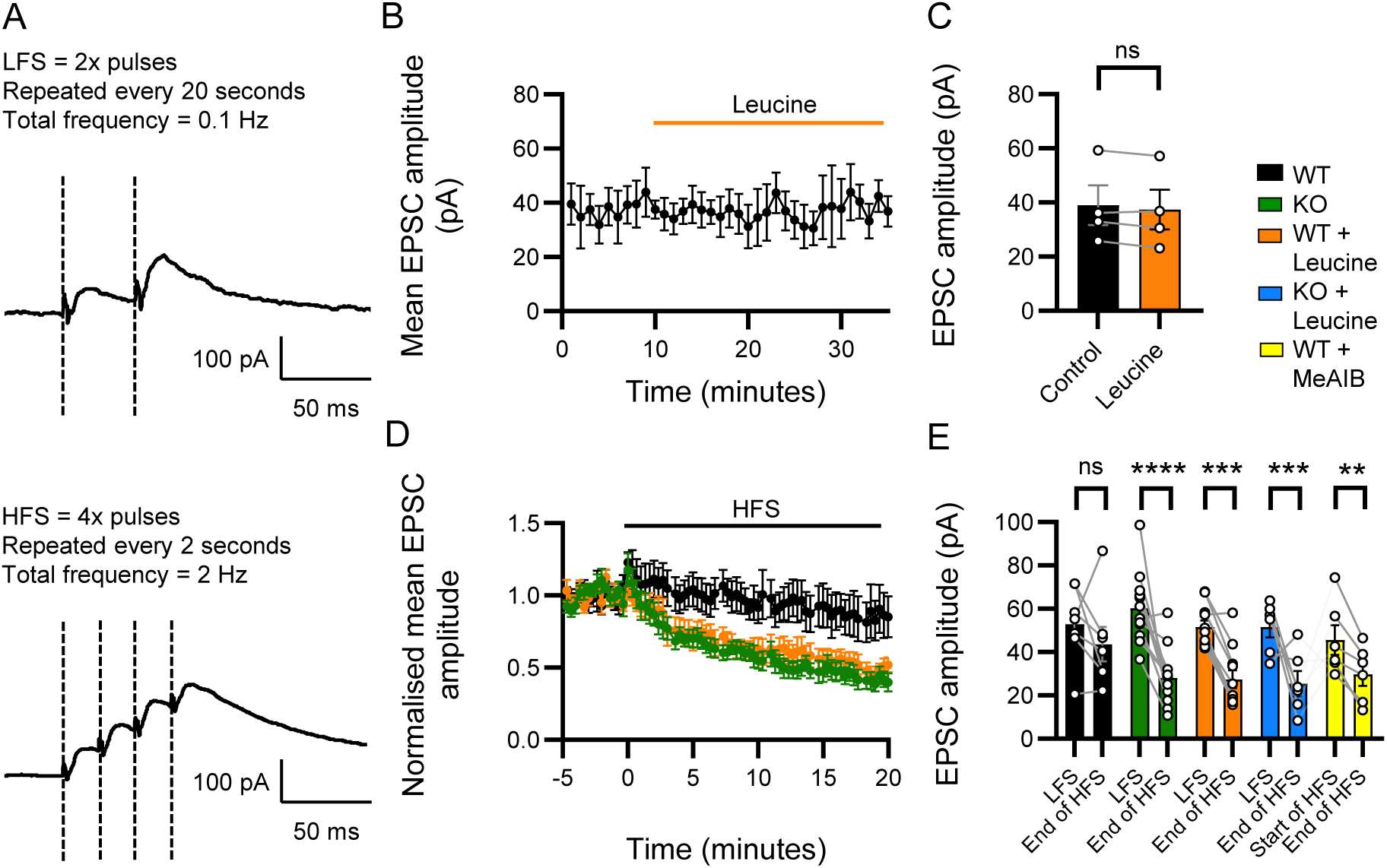
Synaptic transmission is reduced at mossy fibre – CA3 synapses during high frequency stimulation in brain slices when glutamine transport via NTT4 is absent. **(A)** Glutamate release from mossy fibre boutons elicits NMDA-mediated EPSCs in CA3 pyramidal cells when low-frequency (0.1 Hz; LFS; Top) and high frequency (2 Hz; HFS; Bottom) stimulation is applied to presynaptic axons. **(B)** Addition of leucine (10 mM; orange) to the ACSF during LFS does not lead to a reduction in mean EPSC amplitude, suggesting low frequency activity can continue independent of NTT4 over this timeframe. **(C)** Plot showing mean EPSC amplitudes during control (black) and following wash-in of 10 mM leucine (orange). There is no difference in EPSC amplitude during the control period (black) and after addition of 10 mM leucine (orange) when LFS is applied to presynaptic axons. **(D)** HFS does not affect EPSC amplitudes in slices from WT animals (black). In slices treated with leucine (orange) and slices from NTT4 KO animals (green) the EPSC amplitudes reduce, suggesting a depletion of presynaptic glutamate. **(E)** Plot showing mean EPSC amplitudes during LFS and during the last 5 minutes of HFS. Initial EPSC amplitudes in LFS are comparable in the different groups (Figure S5), By the end of HFS, EPSC amplitudes are significantly smaller in WT brain slices treated with leucine (orange) and in KO slices (green) whereas they have not reduced in WT control (black). The addition of leucine to KO slices (blue) causes a reduction of the same magnitude, ruling out the possibility that leucine is affecting the reduction via off-target mechanisms. The addition of 20 mM MeAIB to WT slices (yellow) caused a similar reduction of EPSC amplitude during HFS. Data presented as means ± SEM. Data points in C & E indicate individual mice. ‘ns’ = no significant difference, ‘***’ = p < 0.001 ‘****’ = p < 0.0001.

In contrast to continual LFS, switching to intermittent high frequency stimulation (HFS; 4 pulses 20 ms apart, repeated every 2s, Figure 4A- bottom) is calculated to quickly deplete the presynaptic glutamate pool, necessitating reliable glutamate replenishment to maintain neurotransmission (Marx et al., 2015). Under control conditions with uncompromised glutamine transport, 20 minutes of HFS did not cause a significant reduction in EPSC amplitude (-17.6 %, LFS = 52.9 ± 6.6 pA, v end of HFS = 43.6 ± 8.0 pA, n = 7, p = 0.21, Figure 4D). This demonstrates that high frequency stimulation alone does not appreciably affect the amount of glutamate released by the presynaptic terminals over this timeframe.

In contrast, when NTT4 was absent, EPSC amplitudes reduced by more than half of their original amplitude (-58.3 %, LFS = 60.7 ± 5.1 pA, v end HFS = 25.3 ± 2.9 pA, n = 11, p < 0.0001, Figure 4D) and were significantly smaller than the amplitudes recorded in WT slices during this time (43.6 ± 8.0 pA, n = 7, p < 0.05, Figure 4D & E). Similarly, in WT slices treated with 10 mM leucine, EPSCs also reduced to around half of their original amplitude (-46.8 %, LFS = 51.6 ± 2.9 pA, v HFS = 27.4 ± 4.1 pA, n = 11, p < 0.001, Figure 4D) and were significantly smaller than WT (43.6 ± 8.0 pA, n = 7, p < 0.05) but no different to KO (25.3 ± 2.9 pA, n = 11, p = 0.73, Figure 4D & E). The decrease in EPSC amplitude was not accompanied by change in paired-pulse ratio, showing that it is not the result of a modulation in presynaptic release probability (Figure S2).

Applying leucine to slices from NTT4 KO mice did not cause any additional depletion of NMDA-EPSCs during HFS (KO + leucine = -50.6 %, n = 6, v KO p = 0.99, v WT + leucine p = 0.78, Figures 4E & S3), indicating that the effect of leucine was via inhibition of NTT4. Addition of the amino acid analogue MeAIB to WT slices, which is a broad substrate for both NTT4 and other glutamine transporters, caused a similar reduction in EPSC amplitude from the start to end of HFS (- 34.7 %, from 45.6 ± 6.9 pA to 29.8 ± 5.3 pA, n = 6, p < 0.01, Figures 4E & S4). The similar effects of leucine and MeAIB indicate that other MeAIB sensitive transporters that do not bind leucine (such as SNAT1/2: Varoqui et al., 2000; Yao et al., 2000; Chaudhry et al., 2002b) do not further contribute to maintaining the glutamate supply under these conditions. Taken together, these results show that NTT4 is required to maintain rapid synaptic activity at mossy fibre - CA3 synapses.

### NTT4 is required for sustained synaptic transmission at Schaffer collateral - CA1 synapses

Mossy fibre boutons are large presynaptic terminals that are amenable to patch-clamp studies. However, they may not represent typical brain synapses. Therefore, to investigate whether NTT4 plays the same role at more representative synapses we measured local field potentials at Schaffer collateral - CA1 synapses during LFS (2 pulses at 25 Hz, repeated every 20 s, Figure 5A- top) and HFS (2 pulses at 25 Hz, repeated every 2 s, Figure 5A- bottom). For these experiments we measured AMPA-mediated field EPSCs (fEPSCs) while blocking NMDARs to inhibit plasticity-related changes in glutamate release.

**Figure 5.**
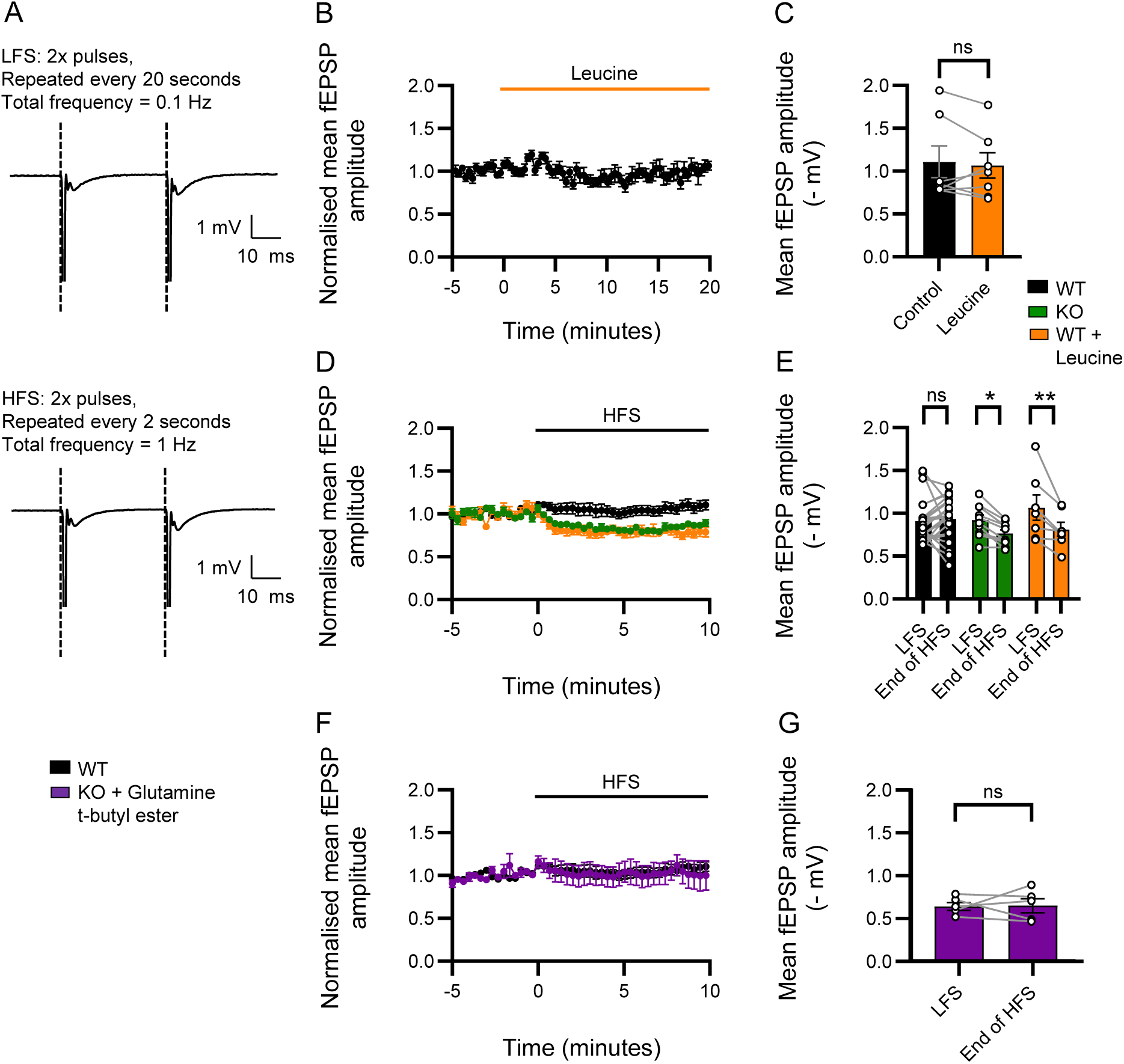
Synaptic transmission is reduced at Schaffer collateral – CA1 synapses during high frequency stimulation in brain slices when glutamine transport via NTT4 is absent. **(A)** Glutamate release from Schaffer collateral presynaptic terminals elicits AMPA-mediated field EPSPs (fEPSPs) in CA1 *stratum radiatum* when low-frequency (0.1 Hz; LFS; Top) and high frequency (1 Hz; HFS; Bottom) stimulation is applied to presynaptic axons. **(B)** Addition of leucine (10 mM; orange) to the ACSF during LFS does not lead to a reduction in mean fEPSP amplitude. suggesting low frequency activity can continue independent of NTT4 over this timeframe. **(C)** Plot showing mean fEPSP amplitudes during control (black) and following wash-in of 1 mM leucine (orange). There is no difference in fEPSP amplitude during the control period (black) and after addition of 10 mM leucine (orange) when LFS is applied to presynaptic axons. **(D)** HFS does not affect fEPSP amplitudes in slices from WT animals (black). In slices treated with leucine (orange) and slices from NTT4 KO animals (green), fEPSP amplitudes reduce, suggesting a depletion of presynaptic glutamate. **(E)** Plot showing mean fEPSP amplitudes during LFS and by the end of HFS in all groups. By the end of HFS, fEPSP amplitudes are significantly smaller in WT brain slices treated with leucine (orange) and in KO slices (green), whereas they have not reduced in WT control (black). **(F)** Incubation of KO slices in glutamine t-butyl ester (purple) prevents the amplitude reduction during HFS, with amplitudes remaining stable as in WT (black). **(G)** Mean amplitudes (mV) of fEPSPs in KO slices incubated in glutamine t-butyl ester during LFS vs the last 5 minutes of HFS show no change. All data presented as means ± SEM. Data points in C, E & G indicate individual recordings. ‘ns’ indicates no significant difference, ‘**’ = p < 0.01.

Similar to the results seen in CA3, the addition of 1 mM leucine to WT slices during LFS did not cause a reduction in fEPSP amplitudes over 20 minutes (from -1.1 ± 0.19 mV to -1.1 ± 0.15 mV, p = 0.56, n = 7, Figure 5B & C). In a separate set of recordings, there were no differences in fEPSP amplitudes during LFS for WT slices (-0.9 ± 0.05 mV, n = 23), KO slices (-0.9 ± 0.06 mV, n = 10) and WT slices treated with 1 mM leucine prior to recording (-1.1 ± 0.15 mV, n = 7), and fEPSP amplitudes in WT slices did not reduce after 5-10 minutes of HFS (+2.9 %, to -0.9 ± 0.05 mV, n = 23, p = 0.58, 5D & E). In contrast, fEPSPs reduced in slices from KO animals (-17.1 %, to -0.8 ± 0.04 mV, n = 10, p < 0.01, Figure 5D & E) and in WT slices treated with leucine (-23.7 %, to -0.8 ± 0.08 mV, n = 7, p < 0.01, Figure 5D & E). The reduction in fEPSC amplitude was not accompanied by a change in paired-pulse ratio (Figure S6), indicating that presynaptic release probability remained unaltered.

To demonstrate that the reduction in EPSC amplitude in NTT4-KO mice was a result of a deficit in intracellular glutamine supply, we reversed this effect by the direct addition of internal glutamine to the presynaptic cells. This was achieved by incubating the slices in 1 mM glutamine t-butyl ester, which enters the cells and is de-esterified to resupply the internal glutamine, independent of NTT4 activity. Accordingly, with the addition of glutamine t-butyl ester incubation to KO slices, fEPSP amplitudes remaining stable after 5-10 minutes of HFS (+1.3 %, from -0.6 ± 0.05 to -0.7 ± 0.08 mV, n = 5, p = 0.93, Figure 5F & G), providing confirmation that the decline in EPSC amplitudes in NTT4 KO animals is due to a depletion of intracellular glutamine.

In addition to fEPSC amplitudes, the results were confirmed by analysis of the fEPSC slopes, with WT recordings remaining stable after 5-10 minutes of HFS (-7.9 %, from 0.42 ± 0.04 to 0.39 ± 0.04 mV/ms, n = 23, p = 0.15) as well as KO slices incubated in glutamine t-butyl ester (-15.9 %, from 0.37 ± 0.04 to 0.31 ± 0.06 mV/ms, n = 5, p = 0.23). In slices from NTT4 KO animals however, there was a reduction in slope (-23 %, from 0.42 ± 0.04 to 0.32 ± 0.03 mV/ms, n = 10, p < 0.01) and there was also a reduction in slope in WT slices treated with leucine (-35.0 %, from 0.47 ± 0.07 to 0.30 ± 0.04 mV/ms, n = 7, p < 0.001).

These findings, similar to the results obtained in whole-cell CA3 recordings, show that NTT4 is also necessary to support ongoing high-frequency synaptic transmission at Schaffer collateral – CA1 synapses by facilitating glutamine entry into the presynaptic terminal.

### Glutamate content of synaptic vesicles is reduced in NTT4 KO mice following HFS

The reduced synaptic transmission seen at both mossy fibre - CA3 and Schaffer collateral - CA1 synapses of NTT4 KO mice following high frequency stimulation was hypothesised to reflect a depletion of presynaptic glutamate and therefore a reduction in presynaptic vesicle glutamate content (Ishikawa et al., 2002). To confirm this, miniature EPSCs were recorded from CA3 pyramidal cells before and after HFS. Due to the need to evoke action potentials during HFS, recordings were not made in the presence of TTX which would typically be used to isolate spontaneous single-vesicle release events. To exclude mEPSCs originating from action potential-induced multi-vesicle release, those with an amplitude of 100 pA or more (approximately 3% of detected events) were omitted from the analysis.

In cells from WT control mice, mEPSC amplitudes were unchanged following HFS (-0.01 %, before HFS = -26.8 ± 1.7 pA vs after HFS = -26.2 ± 1.7 pA, n = 16, p = 0.56, Figure 6A - E). In contrast mEPSC amplitudes in NTT4 KO cells were significantly reduced following HFS (-17.2 %, before = -31.9 ± 4.5 pA vs after = -26.4 ± 3.9 pA, n = 5, p < 0.05. Figure 6A - E). The mean mEPSC amplitudes for each group prior to stimulation was not significantly different (p = 0.11), showing that the reduction in NTT4 cells was not due to pre-existing differences between groups.

**Figure 6.**
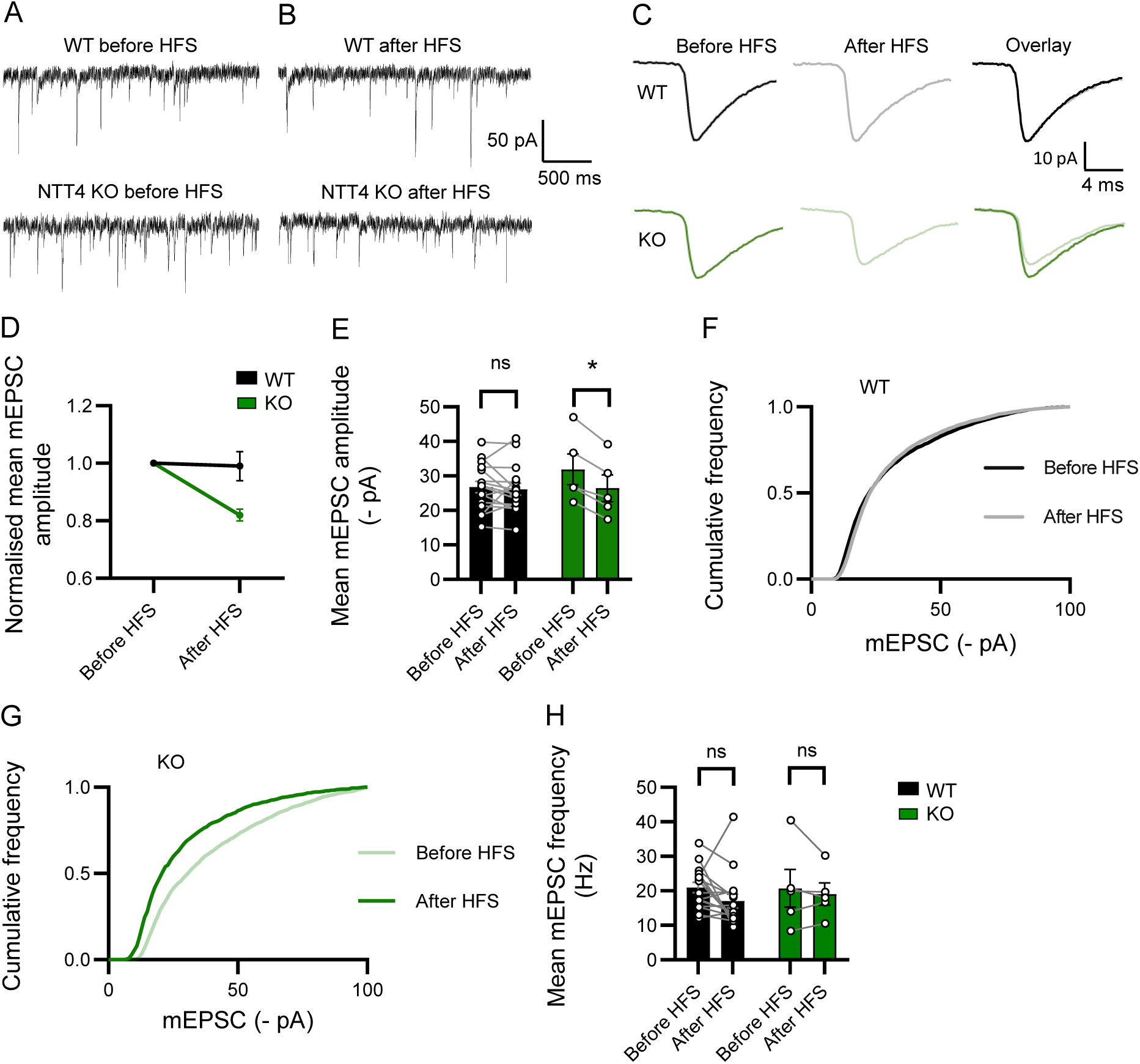
Vesicular glutamate content is reduced following HFS in NTT4 KO mice but not WT control animals. **(A)** Example mEPSC traces showing amplitudes before HFS in WT control and NTT4 KO mice. **(B)** Example mEPSC traces showing amplitudes after HFS in WT control mice and NTT4 KO mice. **(C)** Averaged mEPSCs from one WT (top) and KO (bottom) recording shown in A and B, before (left) and after HFS (middle) and both overlaid (right). **(D)** Normalised mean mEPSC amplitudes highlight the reduction in NTT4 KO mice (green) to around 80 % of their original size (green) whereas WT control (black) do not change. **(E)** Mean mEPSC amplitudes are reduced in NTT4 KO mice (green) but not WT control (black) after HFS, showing that synaptic vesicles contain less glutamate following elevated synaptic activity when NTT4 is absent. Data points show individual recordings. **(F)** Cumulative frequency plot shows the sample distribution for mEPSC amplitudes in WT mice is not changed after HFS (grey line = before HFS, black line = after HFS). **(G)** Cumulative frequency plot for NTT4 KO recordings shows the sample distribution is changed following HFS (light green line = before HFS, dark green line = after HFS), indicating smaller mEPSCs following HFS (p < 0.0001, Kolmogorov-Smirnov test). **(H)** Mean mEPSC frequency is not significantly reduced in either WT control (black) or NTT4 KO (green) cells. Data points show individual recordings. Error bars indicate SEM. ‘ns’ indicates no significant difference, ‘*’ indicates significant difference with p < 0.05.

To compare the sample distributions of mEPSC amplitudes in WT and NTT4 KO cells before and after HFS, a two-sample Kolmogorov-Smirnov test was used. For WT recordings, the maximum difference between the distributions was D = 0.11 (p = 0.59, Figure 6F), whereas for KO, D = 0.32 (p < 0.0001, Figure 6G) showing that the distribution of KO but not WT mEPSC amplitudes are significantly changed following HFS.

The frequency of mEPSCs was unchanged following stimulation in either group (WT before HFS = 21.0 ± 1.5 Hz vs after HFS = 17.0 ± 2.0 Hz, n = 16, p = 0.06, KO before HFS = 20.7 ± 5.4 Hz vs after HFS = 19.0 ± 3.2 Hz, n = 5, p = 0.63, Figure 6H).

This evidence confirms that NTT4 KO mice have reduced glutamate content in presynaptic vesicles following elevated synaptic activity, whereas when NTT4 is intact, glutamate supply is maintained. Further, it supports the conclusion that NTT4 provides glutamine as a glutamate precursor to presynaptic terminals.

### NTT4 is necessary for normal retention of conditioned memory associations

As we have shown that NTT4 KO animals exhibit changes in synaptic transmission in the hippocampus, a region vitally necessary for learning and memory processes, we hypothesised that differences will exist in hippocampus-dependent memory in these animals. We therefore used a trace fear conditioning paradigm which has been shown to assess CA1 and CA3-dependent memory function (Sellami et al., 2018; Al Abed et al., 2020). This protocol involves the delivery of an aversive foot shock (the unconditioned stimulus) paired with a non-aversive tone (the conditioned stimulus) separated by a ‘trace’ period of either 20 or 40 seconds. Healthy adult mice can form and retain an association between conditioned and unconditioned stimulus in both conditions, demonstrating ‘temporal binding’. Temporal binding has been shown to depend on CA1 activation (Huerta et al., 2000; Al Abed et al., 2020). Memory for the context of an event is also hippocampus-dependent, and has been shown to depend on function of CA3 (Kesner and Lee, 2002).

During the conditioning phase which pairs the tone and foot shock, both groups of mice in the 20 and 40-second trace conditions showed increasing fear of the tone with each subsequent pairing (time spent freezing during tone and 20-second trace for WT, at first = 12.5 ± 4.3 %, second = 37.3 ± 4.8 %, p <0.001, and third pairing = 59.5 ± 5.1 % p <0.001, n = 12 and for KO at first = 6.3 ± 2.3 %, second = 23.0 ± 5.8 % p <0.01 and third pairing = 50.5 ± 4.9 %, p <0.001, n = 8, Figure 7B). Time spent freezing during tone and 40-second trace for WT, at first = 14.4 ± 1.8 %, second = 43.2 ± 4.1 %, p <0.001, and third pairing 65.6 ± 6.0 %, p <0.01, n = 12, and for KO at first = 20.0 ± 2.5 %, second = 41.6 ± 3.1 %, p <0.001, and third pairing 55.2 ± 5.7 %, p <0.05, n = 15, Figure 7E). Average total freezing during conditioning was not different between groups (20-second trace, WT = 54.7 ± 6.2 s, n = 12 vs KO = 39.9 ± 3.6 s, n = 8, p = 0.09, 40-second trace, WT = 86.3 ± 6.2 s, n = 12 vs KO = 81.8 ± 4.7 s, n = 15, p = 0.55, Figure 7H), indicating no differences existed in general activity levels.

**Figure 7.**
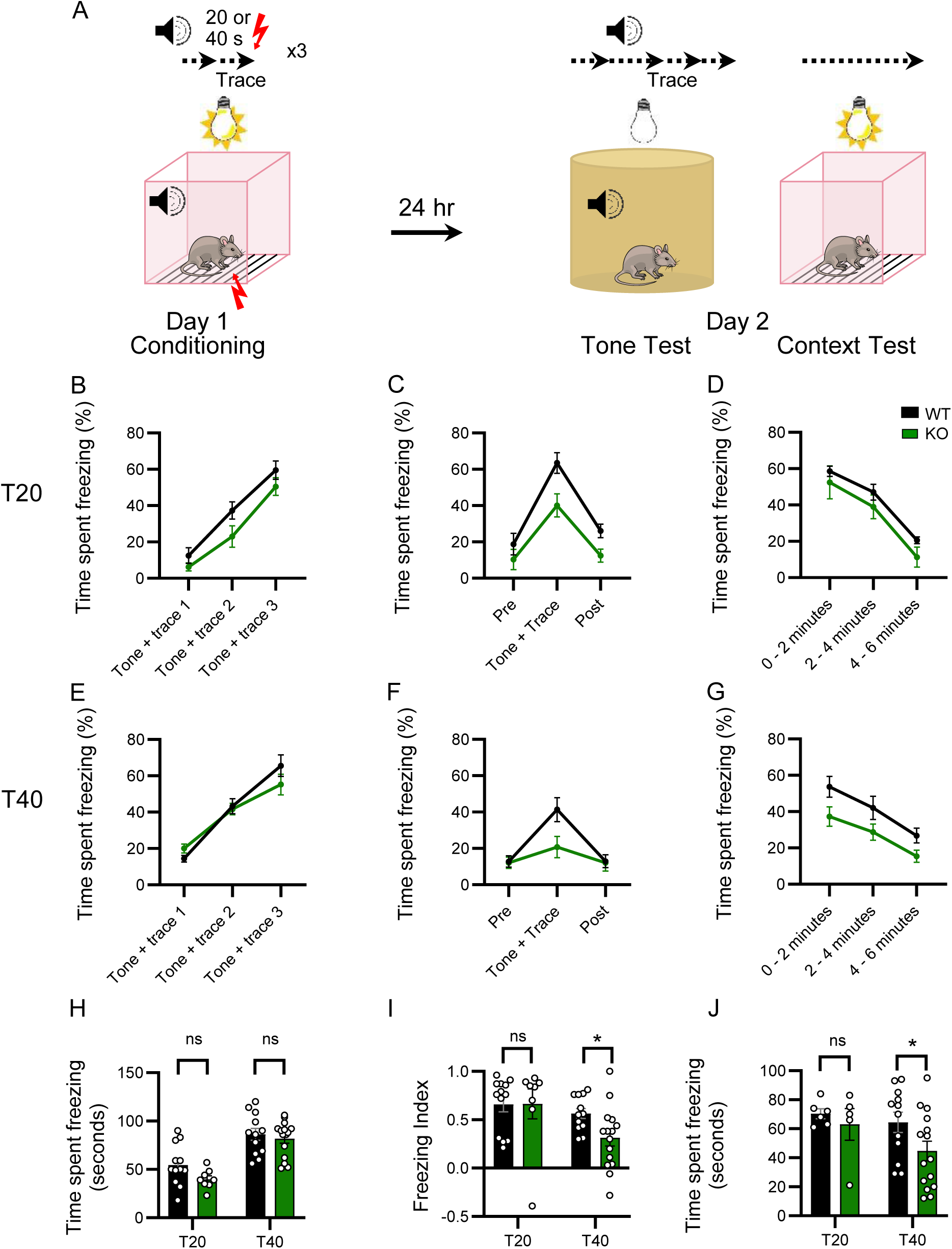
NTT4 KO mice have impaired retention of conditioned memory associations when conditioned and unconditioned stimuli are separated by a longer (40 s), but not a shorter (20 s) temporal gap. **(A)** Schematic diagram of the trace fear conditioning protocol. Mice are placed into a conditioning chamber and a tone is sounded for 30 seconds. A ‘trace’ period of 20- or 40-seconds then elapses before a foot shock (0.2 mA) is delivered. This is repeated 3 times. The following day, mice undergo a tone test, in which the tone is presented while the mouse is in a different chamber, and a context test with no tone, but in the same chamber as the previous day. No foot shock is present during either test. Time spent freezing is measured throughout. **(B)** Comparison of time (%) spent freezing with each pairing of the tone and foot shock in the conditioning phase on day 1 when conditioned with the 20 second trace, WT (black) and NTT4 KO mice (green) both learned to associate the tone and foot shock (indicated by increasing % of time freezing during each subsequent tone and trace period). **(C)** Comparison of time (%) spent freezing for WT (black) and NTT4 KO mice (green) during the pre-tone, tone and trace, and post-tone intervals of the tone test, after being conditioned with the 20 second trace. **(D)** Comparison of time (%) spent freezing for WT (black) and NTT4 KO mice (green) during each 2-minute interval of the context test, after being conditioned with the 20 second trace. **(E - G)** Same as B – C but for mice conditioned with the 40 second trace. **(H)** During the conditioning phase with both the 20 and 40 second trace, WT (black) and NTT4 KO mice (green) spent an equal amount of time freezing. **(I)** During the tone test on day 2, NTT4 KO mice (green) spent less time freezing during the tone and trace relative to the pre-tone interval, than the WT mice (black), when conditioned with the 40 second but not the 20 second trace. **(J)** Likewise, during the context test, NTT4 KO mice (green) that had been conditioned with the 40 second, but not the 20 second trace, spent less time freezing in the first 2 minutes than the WT mice (black). Data presented as means ± SEM. Data points in H-J indicate individual mice. ‘ns’ indicates no significant difference, ‘*’ indicates p < 0.05.

There were no differences between groups in the subsequent tests when animals were conditioned with a 20-second trace period (In the tone test, freezing index for WT = 0.65 ± 0.08, n = 12 vs KO = 0.66 ± 0.16, n = 8, p = 0.91, Figure 7C & I, and in the context test, total freezing in the first two minutes for WT = 71.8 ± 3.8 s, n = 5 vs KO = 63.0 ± 10.9 s, n = 5, p = 0.56, Figure 7D & J). For those conditioned with a 40-second trace, NTT4 KO animals performed significantly worse than WT controls during the tone test (freezing index for WT = 0.56 ± 0.05, n = 12 vs KO = 0.31 ± 0.08, n = 15, p < 0.05, Figure 7F & I), indicating reduced fear memory. Similarly, NTT4 KO mice initially performed worse than WT controls when re-exposed to the original conditioning chamber (freezing for 64.4 ± 6.9 s, n = 12 vs KO = 44.7 ± 6.7 s, n = 15, p < 0.05, Figure 7D).

This indicates that not only does NTT4 play a role in maintaining presynaptic glutamate for normal synaptic transmission, it also plays a measurable role in normal hippocampus-dependent behaviours such as memory retention.

### Nest building, anxiety behaviour and social preference are altered by NTT4 deletion

Given the demonstrated importance of NTT4 in normal memory retention, we reasoned that the behavioural effects of NTT4 deletion were likely to be widespread. Indeed, in the human cohort studied by Iqbal et al. (2015) the behavioural effects of *SLC6A17* mutation extended beyond intellectual disability to impaired social, locomotor and mood – related behaviours. Nest building is fundamental mouse behaviour for providing shelter and thermoregulation. The construction of nests is straightforward to assess and a commonly used measure of impairment in mouse models of pathology including intellectual disability (Heller et al., 2014; Kaur et al., 2014; Yuan et al., 2018) and Alzheimer’s disease (Wesson and Wilson, 2011). Using an established scoring system (Figure 8A) nests were rated 0 – 5. While WT mice were proficient in nest building ability (nest score = 3.4 ± 0.3, n = 9, Figure 8B), NTT4 KO mice showed a considerable impairment (nest score = 0.9 ± 0.3, n = 18, Mann-Whitney U test p < 0.0001, Figure 8B). This finding highlights a role of NTT4 in supporting activities of daily living.

**Figure 8.**
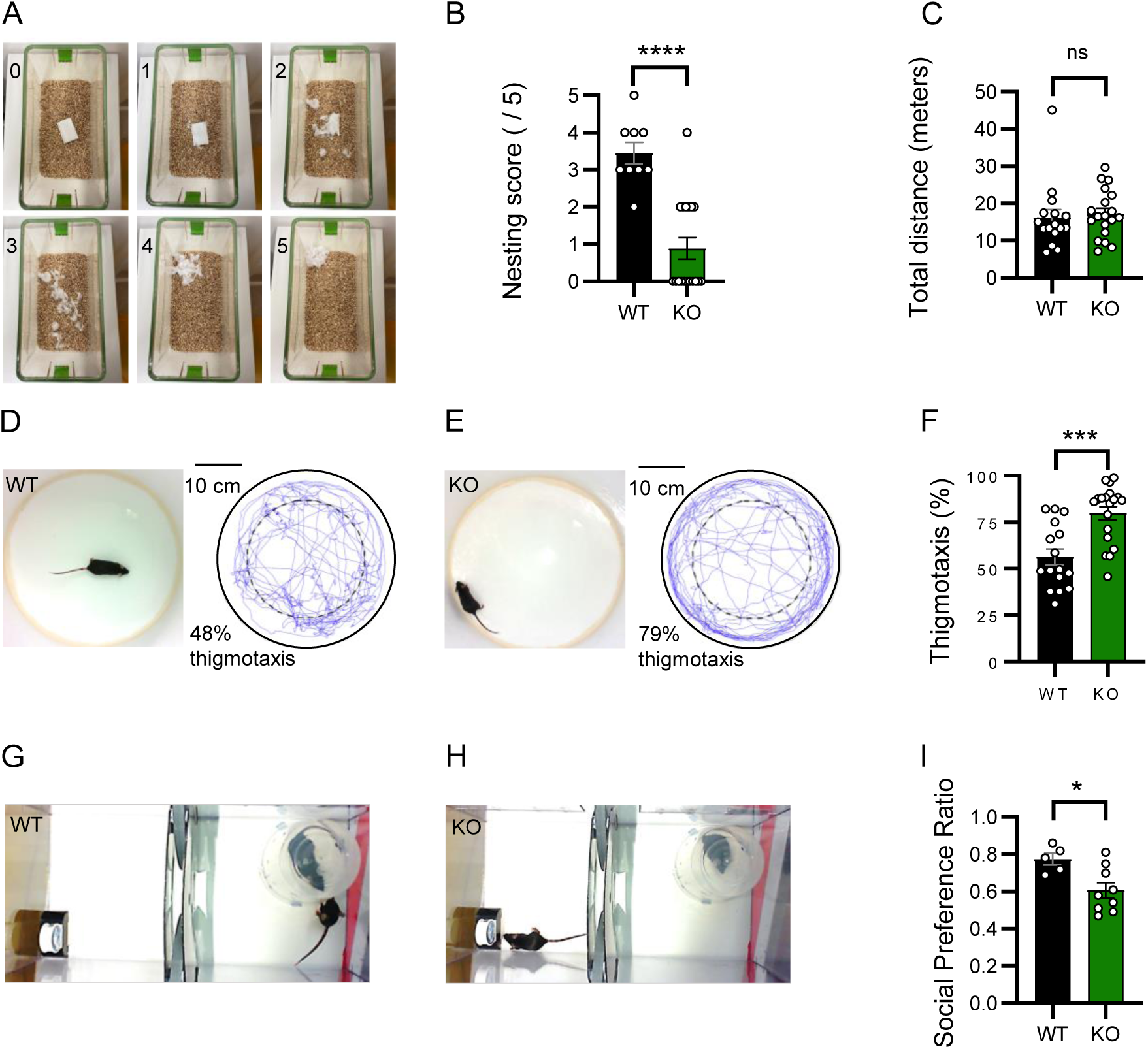
NTT4 KO mice show deficits in activities of daily living, increased anxiety and reduced social preference. **(A)** Example of the rating scale used to score the incorporation of nestlet material into a nest after being left undisturbed for 24 hours. ‘0’ shows completely intact nestlet and ‘5’ shows complete incorporation of nestlet into a nest comprising a cup-shaped base and domed walls. **(B)** NTT4 KO mice (green) scored lower average scores than WT mice (black) on the nesting scale depicted in (A). The same results were observed in both the male and female cohort of mice (Figure S7). **(C)** In an open field test, NTT4 KO mice (green) and WT (black) mice traversed similar average distances over 4 minutes of free exploration. **(D)** Example screenshot (left) from open field test shows a WT mouse exploring the centre of arena. The movements of this mouse during 4 minutes of free exploration (blue trace) shows approximately half of the movements were around the outer edge (48 % thigmotaxis, which was close to the group mean of 56%.) and an approximately equal proportion in the centre of the arena. **(E)** As in (D) however, screenshot (left) from open field test shows an NTT4 KO mouse displaying thigmotaxis; the tendency to remain in the outer perimeter of the arena. The movements of this mouse during 4 minutes of free exploration (blue trace), shows a high proportion of thigmotaxis (79 %), representative of the group mean. **(F)** NTT4 KO mice (green) showed greater thigmotaxis than WT mice (black) over 4 minutes in the open field test. **(G)** A screenshot from the two-chamber social preference test shows a WT mouse interacting with the novel mouse, contained within the right chamber. **(H)** As in (G), but here, the screenshot shows an NTT4 KO mouse inspecting the novel object in the left chamber **(I)** NTT4 KO mice (green) showed a smaller social preference ratio than WT mice (black), calculated as time spent interacting with the novel mouse / the time spent interacting with the novel mouse and the novel object (mouse / mouse + object) during 5 minutes of the test, indicating they had a reduced preference for social interaction. Data presented as means ± SEM. Data points in B, C, F & I indicate individual mice. ‘ns’ = no significant difference, ‘*’ = p < 0.05, ‘***’ = p < 0.001 ‘****’ = p < 0.0001.

As NTT4 mutation has previously been shown to be associated with disordered mood in humans, another area where behavioural impairment in NTT4 KO mice might be evident is anxiety behaviours. Mood and anxiety disorders, whilst distinct, are highly comorbid and anxiety in mice can be tested more directly than disordered mood. As a prey species, mice are generally hesitant to explore open spaces, and as such they display thigmotaxis; the tendency to restrict movements to the area around the perimeter of an open arena, before venturing into the centre. Accordingly, mice with heightened anxiety have been shown to display more thigmotaxis and less crossings into the centre of the arena (Treit and Fundytus, 1988; Simon et al., 1994). In an open field test where mice were allowed to explore freely for 4 minutes, WT and NTT4 KO mice traversed a similar total distance (WT = 16.1 ± 2.2 m, n = 16 vs KO = 17.2 ± 1.5 m, n = 19, p = 0.69, Figure 8C). However, NTT4 KO mice showed more thigmotaxis (79.9 ± 5.6 %, n = 16), than WT animals (56.2 ± 4.1 %, n = 19, p < 0.0001, Figures 8D - F) suggesting they are more anxious than healthy controls.

Social preference is another behavioural domain linked to both *SLC6A17* function in humans (Iqbal et al., 2015) and glutamatergic neurotransmission in mice (Adamczyk et al., 2012; Li et al., 2023). As mice are social animals they typically choose to interact with a novel mouse in preference to a novel object. In a two-chamber social preference test, WT mice spent 77 % of the time engaged in social interaction (0.77 ± 0.03, n = 5, Figures 8G & I). In contrast, NTT4 KO mice showed a reduced preference for social interaction, spending only 61 % of their time interacting with a novel mouse (0.61 ± 0.04, n = 9, p < 0.05, Figures 8H & I). This confirms that NTT4 plays a role in normal social behaviour in mice, and in its absence, the preference for social interaction is reduced.

Taken together this set of results demonstrate the importance of NTT4 in regulating glutamate production via the glutamate-glutamine cycle and highlight its role in controlling levels of excitatory neurotransmission and thus influencing animal behaviour.

## Discussion

Our results clearly demonstrate that NTT4 is a glutamine transporter, which is functional in the presynaptic plasma membrane of glutamatergic synapses. It controls the levels of glutamate release via sequestration of presynaptic glutamine and is thus a key component of the glutamate-glutamine cycle. Modulation of this transporter by genetic knockout or acute pharmacological inhibition reveal that NTT4, and hence the glutamate-glutamine cycle, is required to maintain neurotransmitter levels during bursts of high-demand synaptic transmission. Consistent with this impairment of synaptic function, mice lacking NTT4 have deficiencies in hippocampus-dependent behaviours such as long-term memory processing, as well as reduced engagement in activities of daily living and sociability, and an increase in anxiety behaviour.

Previous evidence has shown that NTT4 is mainly located in the membrane of small synaptic vesicles (Fischer et al., 1999; Masson et al., 1999; Parra et al., 2008; Jia et al., 2023), with a suggestion that its physiological role is to load vesicles with glutamine (Jia et al., 2023). However, counter to this view, our direct patch-clamp recordings demonstrate that NTT4 is functional in the nerve terminal plasma membrane of intact tissue, where it transports glutamine into the presynaptic cytoplasm. This evidence of plasma membrane location is further supported by our data showing NTT4-mediated uptake of glutamine into synaptosomes. These apparently contradictory findings concerning NTT4’s location can be partially reconciled by our observations of presynaptic glutamine transport in the calyx of Held presynaptic terminal (Billups et al., 2013). This study shows that presynaptic glutamine transporters are located on vesicles in the nerve terminal and are repeatedly trafficked to and from the plasma membrane when the terminal is active or quiescent. As insertion is SNARE-dependent and removal is clathrin-dependent (Billups et al., 2013), this suggests that there is store of glutamine transporters on vesicles that are supplied to the plasma membrane, as required by the demand for neurotransmitter replenishment.

In addition to glutamine, NTT4 can transport other neutral amino acids, including alanine, proline, glycine and leucine (Parra et al., 2008; Zaia and Reimer, 2009). While glutamine is the most abundant amino acid in the brain extracellular fluid, with CSF concentrations approximately 10 to 100-fold higher than the other NTT4 substrates (Reichel et al., 1995), the transporter’s physiological role may also involve the sequestration of different amino acids. Nevertheless, our ^13^C NMR spectroscopy and LC-MS data clearly show that tissue glutamine levels are reduced when NTT4 is absent. Additionally, the fact that we can reverse the effects of NTT4 depletion on glutamatergic neurotransmission by the replenishment of intracellular glutamine using application of a glutamine ester, strongly suggests that NTT4 exerts its main physiological role via the transport of glutamine, rather than an alternative substrate.

Our direct recordings from mossy fibre boutons demonstrate that the presynaptic glutamine transporter currents are entirely mediated by NTT4. In the absence of NTT4 other glutamine transporters, such as B^0^AT2 (*SLC5A15*) or SNAT1 (*SLC38A1*), do not contribute to the glutamine influx, at least at this synapse. Previous studies have suggested that the system A transporters e.g. SNAT1 (*SLC38A1*) perform the role of presynaptic glutamine transport (Armano et al., 2002; Chaudhry et al., 2002a), although localisation studies do not find these transporters in synaptic terminals (Mackenzie et al., 2003; Melone et al., 2006). Discrepancies in the results around system A transporters’ involvement in excitatory transmission may arise from the use of MeAIB as substrate inhibitor (Christensen et al., 1965; Reimer et al., 2000; Chaudhry et al., 2002b). While it is frequently used as a system A inhibitor, MeAIB is non-specific and also interacts with NTT4 (Zaia and Reimer, 2009), and other neuronal glutamine transporters (Erickson, 2017). Studies that have ascribed a physiological role for system A in the brain based on the effects of MeAIB (e.g. in epileptiform activity: Bacci et al., 2002; Tani et al., 2010) may therefore have overlooked the role of alternative MeAIB-sensitive transporters, such as NTT4. Accordingly, recordings of glutamine transporters at the calyx of Held presynaptic terminal (Billups et al., 2013) exhibit MeAIB sensitivity, which is consistent with presynaptic NTT4 at this synapse also.

Given the localisation of NTT4 in the presynaptic plasma membrane and the evidence showing that it imports glutamine, it is likely that a significant portion of the presynaptic glutamate store is derived from glutamine transported into presynaptic terminals by NTT4. We have shown this by examining metabolites of ^13^C-enriched glucose and acetate through neurons and astrocytes. Mice lacking NTT4 have reduced glutamate and glutamine production in cortex and hippocampus, consistent with impaired presynaptic glutamine transport which would both deplete the levels of presynaptic glutamate deriving from astrocytic glutamine, and in turn, reduce the amount of glutamine originating from released glutamate.

A reduction in the presynaptic glutamate supply has been shown to limit filling of synaptic vesicles at glutamatergic terminals, causing a reduction in levels of excitatory neurotransmission (Ishikawa et al., 2002; Edwards, 2007). Indeed, in brain slices from NTT4-KO mice we observe a reduction of approximately half in EPSC amplitudes in CA3 pyramidal cells when presynaptic axons were stimulated at a high frequency over 20 minutes, due to a presynaptic depletion of glutamate. The same was true for WT slices when NTT4 was acutely inhibited with leucine or MeAIB. In contrast, in WT control slices, high frequency stimulation alone was not enough to cause glutamate to deplete. This reduction in excitatory neurotransmission was observed when examining glutamate release by measuring NMDA-mediated responses at mossy fibre-CA3 synapses (with AMPA receptors blocked) and also by measuring AMPA receptor mediated responses at Schaffer collateral – CA1 synapses (with NMDA receptors blocked), showing that NTT4 is necessary for maintaining presynaptic glutamate supply and adequate levels of synaptic glutamate release during intense activity. This is further supported by the observed reduction in miniature EPSC amplitudes following high-frequency stimulation when NTT4 was absent, showing a reduced vesicular glutamate content. This is consistent with previous data showing that inhibition of the glutamate-glutamine cycle reduces miniature EPSC amplitude during periods of sustained synaptic stimulation (Billups et al., 2013; Tani et al., 2014).

Finally, as it has been shown that *Slc6a17* mutations have significant behavioural effects in humans (Iqbal et al., 2015), we investigated the impact of NTT4 deletion on mouse behaviour and memory *in vivo*. Removal of NTT4 was shown to inhibit memory formation, increase anxiety, impair nest building and reduce the desire for social interaction. For episodic memory in the temporal domain, which is known to be dependent on hippocampal CA1 activation (Huerta et al., 2000), when a 20-second trace period separated the unconditioned and conditioned stimuli, temporal binding was unaffected in NTT4 KO animals and they performed on par with WT controls. In contrast, when the trace period was extended to 40 seconds, NTT4 KO mice, but not WT animals showed a significant reduction in their fear responses to both the conditioned tone and the environmental context, showing impaired temporal binding over this extended gap. A similar effect has previously been shown in ageing mice, where temporal binding over the 40-second timeframe was impaired, but subsequently rescued by optogenetic activation of hippocampal CA1 during the trace interval. Likewise, the conditioned association was impaired in younger animals by optogenetic inhibition of CA1 during the trace (Sellami et al., 2017; Al Abed et al., 2020). Other research suggests that sustained activation is less important for fear memory retention than the activation pattern of fear-related circuits in CA1 (Ahmed et al., 2020). Our findings show that CA1 activation, is impaired by the genetic deletion of NTT4, emphasizing the need for high-fidelity glutamatergic responses in this area. Our results therefore illustrate that memory retention is impaired in NTT4 knockout animals, with mechanisms for this hinted at in past research, such as by Gibbs et al. (1996) who found that inactivation of the glutamine producing enzyme glutamine synthetase impaired memory consolidation in one day-old chicks. Similarly, Cheung et al. (2022) showed that interrupting hemichannel-mediated glutamine efflux from astrocytes impaired memory in a novel object recognition task. These findings support our results, as a similar outcome would be expected if NTT4 is required to supply astrocytic glutamine to the presynaptic terminals. In all cases, glutamine entering the presynaptic terminals would be reduced, leading to a smaller quantity of glutamate being produced and released. Thus, the memory effects seen in our experiments are likely due to impaired encoding resulting from reduced glutamate signalling in NTT4 KO animals.

While this study undoubtedly identifies NTT4 as a key component of the glutamate-glutamine cycle and highlights its involvement in glutamate replenishment, it is clear that a significant proportion of glutamate released from excitatory nerve terminals does not originate from this pathway. Other mechanisms of glutamate recycling or replenishment must exist as a) up to 50% of the glutamate transported into astrocytes is metabolised in the astrocytic TCA cycle and is not used to produce glutamine (McKenna, 2007); b) NTT4-KO mice have a relatively subtle phenotype, and survive well into adulthood; and c) removal of NTT4 has little effect on basal levels of synaptic transmission, and is only partially inhibitory when the need for glutamate replenishment is enhanced, during periods of high demand. It is therefore interesting to speculate that pharmacological inhibition of NTT4 could have clinical effects by reducing excessive excitatory neurotransmission e.g. during seizures, while leaving moderate transmission intact. We also do not know how long NTT4 remains at the cell surface after fusion of vesicles with the membrane. Elucidating the trafficking process in more detail, and its control mechanisms, would be an exciting avenue for future research.

In summary, we provide evidence that shows NTT4 is responsible for supplying presynaptic terminals with an appreciable portion of glutamate precursor, and demonstrate that the glutamate-glutamine cycle is necessary for sustaining rapid ongoing synaptic transmission. Further, this transporter and its glutamate replenishment play a key role in normal memory formation and other hippocampus-relevant behaviours.

## Materials and Methods

### Animals

All experiments were conducted with the approval of the Australian National University Animal Experimentation Ethics Committee under the National Health and Medical Research council code of practice (A2014/58, A2014/59, A2017/49, A2020/38, A2021/43 and A2024/331). *Slc6a17^-/-^* (KO) mice in a C57BL/6N background strain (Charles River) were generated within the Australian Phenomics Facility at the Australian National University using the CRISPR/Cas9 system. Slc6a17 guide RNA (gRNAs) sgRNA1 5’-CGAGTCAGTGGCTGACTTGC-3’ sgRNA2 5’-CTCGGTGTTGGCCACCCTCG –3’ and sgRNA3 5’-CAGCGTCATCATGACTGTTA-3’ respectively targeting the exon2, exon7 and exon 8 and Cas9 proteins were obtained from Integrated DNA Technologies. The procedure for mouse pronuclear injection and mouse genotyping was previously described (Waters et al., 2021). The founder retained for downstream analysis exhibited one base pair indels in exon 8 confirmed by Sanger sequencing causing a frame-shift mutation at *Slc6a17*. This mutation scrambles the protein from 411T and prevents cell-surface expression of a functional transporter. Mice of both sexes were used. For electrophysiological recordings mice were aged between postnatal day (P) 21 and 25. For all other experiments, adult mice aged 2 - 6 months were utilised.

### Antibodies

NTT4 and B^0^AT2 (*Slc6a15*) custom antibodies were raised in rabbit by Pineda Antibody services (Germany), against the N-terminus of the relevant proteins. NTT4 antibody was raised against the rat protein and B^0^AT2 antibody against the mouse protein. The rat NTT4 N-terminus was amplified by PCR using primers: rNTT4-NTerm-s C<u>GAATTC</u>ATGCCGAAGAACAGCAAG and rNTT4-NTerm-a C<u>GAATTC</u>TGCAGCTTGCTGTTCCA. *BamHI* and *EcoRI* restriction enzyme sites were added to the sense and antisense primers respectively to assist in insertion into pGEX-2T for the production of recombinant GST fusion protein in *E.coli* BL-21. The mouse B^0^AT2 N-terminus was amplified using primers B^0^AT2-NTerm-s GCGAATTCATATGCCTAAGAATAGCAAAGTG and mB^0^AT2-NTerm-a GCGAATTCTTGCAGCTTACTGTTCCAGGC and inserted into pGEX-2T using the *EcoRI* restriction site.

For the production of the GST fusion proteins, a 1L culture was grown at 28°C in unbaffled flasks and induced with 100 μl of 1 M IPTG when the OD_600_ of the culture was between 0.4 and 0.6. The flasks were shaken for another 5 hours, after which the cultures were collected by centrifugation (5000 g, 10 min), frozen in liquid nitrogen and stored at - 80°C. To release the protein, frozen pellets were thawed on ice, dissolved in 5.0 ml PBS pH 7.4. and sonicated using a MSE 100 Watt Ultrasonic Disintegrator (MSE Technology Applications, USA). Triton-X-100 (1 %) was added to the sonicated cells and the mixture was incubated on ice for 15 minutes. The insoluble debris was removed by centrifugation at 12000 g for 5 minutes at 4 ^0^C in a SS-34 rotor in a Sorvall RC 5C Plus centrifuge. The supernatant was retained and supplemented with Complete EDTA-Free Protease Inhibitor. The protein was bound to Glutathione-Sepharose 4B (Amersham Biosciences, USA) overnight in a rotating incubator at 4°C. On the next day the Sepharose was filled into a column and the protein was eluted using 10 mM reduced glutathione in 50 mM Tris-HCl pH8. The purified protein was used for antibody production.

Other antibodies were purchased from suppliers, including EAAT1 (*Slc1a3*) antibody from Abcam, actin and synaptophysin from Cell Signaling, anti-rabbit and anti-mouse IgG from GE Healthcare and anti-rabbit Alexa 555 from Life Technologies.

### Synaptosomes

Synaptosomes were prepared from mouse cortical tissue following a method described by López-Pérez (1994). Briefly, cerebral cortices (n = 4 – 7 mice per preparation) were differentially centrifuged in a continuous sucrose gradient then purified with an aqueous two-phase system using dextran 500 and polyethylene glycol (PEG) 4000. After purification, synaptosomes were suspended at a 0.5 mg/ml final concentration, in buffer (0.32 M sorbitol, 0.1 mM potassium EDTA and 5 mM potassium phosphate, pH 7.4). Following snap freezing in liquid nitrogen, the suspension was stored at -160°C. Suspensions were thawed on the day of experiment (within 2 weeks of preparation) and kept on ice during use.

### Western blot

25μL of synaptosome preparation (containing 12.5μg of the protein) was mixed with 4 x LDS sample buffer (Invitrogen) and 10 x reducing agent (Invitrogen) followed by incubation at 70°C for 10 min. Samples were separated on a 4-12% Bis-Tris NuPAGE gel (Invitrogen) at 200 V for 1 hour, before blotting onto nitrocellulose membranes (GE Healthcare) for 2 hours at 100 V using a BioRad Mini Trans blot cell. Prestained SeaBlue plus 2 (Invitrogen) was used as a protein standard. Overnight incubation of the nitrocellulose in 5 % milk powder in phosphate-buffered saline (PBS) / Tween (0.1 %) to prevent nonspecific binding was followed by incubation with the primary antibody (NTT4 1:3000, B^0^AT2 1:2000, EAAT1 1:3000, Actin 1:5000, Synaptophysin 1:1000) and 1.5 % milk powder in PBS / Tween (0.1 %) overnight. Blots were washed 3 times with PBS / Tween (0.1 %) then incubated for 4 hours with the secondary antibody in 1.5 % milk powder in PBS / Tween (0.1 %) (anti-rabbit IgG 1:2000-1:5000, and for actin anti-mouse IgG 1:5000) before a final 3 washes. Pierce Ultra HRP was used for detection.

### Amino acid uptake

Synaptosome preparation containing 8-10 µg protein was made up to a total volume of 200 µl with Hank’s buffer (1.26 mM CaCl_2_, 5.4 mM KCl, 0.4 mM KH_2_PO_4_, 0.5 mM MgCl_2_, 0.4 mM MgSO_4_, 136.6 mM NaCl, 10 mM HEPES, 2.7 mM Na_2_HPO_4_, pH 7.5). L-glutamate, L-glutamine, L-proline or L-leucine (90 µM) was added to the preparation along with 0.1 µCi of the same ^14^C labelled amino acid for tracing. Preparations were then left at room temperature for 6 minutes to allow synaptosomes to take up the substrate. The addition of 2 mL ice-cold buffer halted the uptake at the end of this time and the synaptosomes were collected on nitrocellulose filters (0.45 μm). Radioactivity in the isolated synaptosomes was quantified by scintillation counting. Uptake was normalised to the glutamate transport for the corresponding synaptosome preparation.

### Immunohistochemistry

Adult *Slc6a17^-/-^* and ‘wild-type’ C57BL/6N mice were anaesthetised with isoflurane then perfused transcardially with 0.01 M PBS followed by 4 % paraformaldehyde (PFA). Animals were then decapitated and brains removed and postfixed in 4 % PFA until the following day when slices were prepared. Coronal slices 50 µm thick were cut and placed into cryoprotectant (ethylene glycol) then stored at -20°C. Upon removal from storage slices were placed into a 24-well plate and washed twice with PBS to remove cryoprotectant then permeabilised with 0.25 % Triton in PBS. To prevent non-specific binding, blocking solution containing 10 % donkey serum and 0.25 % Triton X-100 in PBS was added to the slices and left for 1.5 hours. Primary antibodies (1:500 NTT4 or B^0^AT2 in blocking solution) were then added the slices and left overnight at 4°C before being brought to room temperature over 2 hours. After 3 x 10-minute washes with PBS, secondary antibody (1:200 anti-rabbit Alexa 555) was added and left for 2 hours in the dark. After 3 further washes, 5 µM DAPI in PBS was added and left for 5 minutes before washing off the slices and mounting them on Livingstone slides with Mowiol and Thermofisher coverslips. Imaging was performed with a Nikon A1 confocal fluorescent microscope (20x, 40x and 63x objectives).

### Preparation of samples for carbon 13 tracing using NMR spectroscopy and LC-MS

Adult wild-type and *Slc6a17^-/-^* mice were injected subcutaneously with 10 µL / g [1- ^13^C] D-glucose (0.3 M) and [1,2-^13^C] acetate (0.6 M) 15 minutes prior to decapitation without anaesthetic (Nilsen et al., 2013). Brains were rapidly dissected with cortex and hippocampi removed, placed in microcentrifuge tubes and snap frozen by submerging tubes in liquid nitrogen. Samples were stored at -80°C until further use. Nuclear magnetic resonance (NMR) spectroscopy was then used to determine the fate of sequestered molecules in the cortical samples, and liquid chromatography-mass spectrometry (LC-MS) to assess metabolites in hippocampal samples.

### Carbon 13 NMR spectroscopy analysis of glutamate and glutamine in cortex

Frozen tissue samples were extracted for ^1^H NMR analysis as described below. Lyophilised extracts were resuspended in ^2^H_2_O containing 2 mM [^13^C] formate as an internal concentration reference and dispensed into 3 mm Norell NMR tubes for analysis. ^1^H, [^13^C]-decoupled ^1^H and [^1^H]-decoupled ^13^C NMR spectra were acquired from the mouse cortex extracts using a Bruker AVANCE III HD 600 NMR spectrometer equipped with a TCI cryoprobe and refrigerated sample changer (Bruker Biospin, Ettlingen Germany), as described previously (Das et al., 2020). Fully relaxed (TR = 30s) ^1^H NMR spectra, with and without decoupling (Rae and Balcar, 2014) were acquired across 24k data points and represented the sum of 32 transients. Acquisition of [^1^H]-decoupled ^13^C NMR spectra (TR = 4s) was across 64k data points and represented the sum of 4096 transients.

Peak areas, integrated in TopSpin 3.6 (Bruker Biospin, Ettlingen Germany) were compared to that of the internal standard [^13^C] formate. In the case of ^13^C spectra, these were adjusted for relaxation and nuclear Overhauser effect by reference to the integrals of standard, fully relaxed (TR = 90s) spectra acquired separately (Rae et al., 2000; Rae and Balcar, 2014). Values were also adjusted relative to the total protein concentration of the sample which was measured from the pellet of each sample following extraction using the method of Lowry (Lowry et al., 1951).

### LC-MS quantification of isotopically enriched glutamate and glutamine in hippocampus

Metabolites were extracted by adding 60 % (v/v) methanol to each sample up to a final volume of 600 μl. To account for extraction losses, [^13^C-5] glutamine (CLM-1822-H-0.1; Cambridge Isotope Laboratories) was spiked into each sample at a final concentration of 15 μM. Each sample was then homogenised using a PRO200 homogenizer (PRO Scientific Inc.) for ten seconds. The homogenates were then snap frozen in liquid nitrogen and kept on dry ice. 200 μl of chloroform was added to each tube and the mixture vortexed for 5 minutes, followed by 5 minutes of centrifugation at 13 000 rpm. 100 μl of the aqueous phase from each sample was the aliquoted into two clean microcentrifuge tubes, which were dried in a vacuum concentrator for three hours.

Each analyte was quantified using one of two analytical methods. For the quantification of amino acids, dried metabolites were solved in 60 μl of 10 mM ammonium acetate : acetonitrile (1 : 9; with 0.15% formic acid) containing 10 μM of [^13^C-U] [^15^N-U] labelled internal standards (MSK-CAA-1; Cambridge Isotope Laboratories). Analytes were separated using a SeQuant ZIC cHILIC 3μm 100Å 150 x 2.1 mm column (EMD Millipore) installed in an Ultimate 3000 RS UHPLC system (Dionex) that was coupled to an Orbitrap Q Exactive mass spectrometer (Thermo Scientific). An increasing ratio of mobile phase A (10 mM ammonium acetate) to mobile phase B (acetonitrile), both containing 0.15% formic acid was used to separate amino acids. The mass filter was set to 73-230 m/z and analytes were ionized in positive mode. (Gauthier-Coles et al., 2021).

For the quantification of GABA, dried samples were incubated with 120 μl of n-butanol containing 3 N acetyl chloride, vortexed for five minutes and incubated at 65°C for 25 minutes. Samples were once again dried in a vacuum concentrator and resuspended with ammonium acetate : acetonitrile (93:7; with 0.15% formic acid) and internal standards at a final concentration of 10 μM. Internal standards were prepared by derivatising citrate and GABA standards in the same manner as the samples, but with non-deuterated n-butanol (615099; Sigma). Analytes were separated using a Kinetex 1.7 μm C18 100Å 100 x 2.1 mm column installed in the same LCMS system with similar parameters as described above but with a few modifications.

In both analyses, the ratio of each analyte and their respective internal standard was calculated and measured against a calibration curve produced from injecting external standards at the beginning of each sequence. Quantities were then adjusted according to the extraction efficiency of each sample, as measured by quantifying the [^13^C-5] glutamine that was originally spiked in prior to sample homogenization. Analyte quantities are normalized to the mass of each hippocampi sample and expressed as picomoles per mg tissue. Blanks and quality controls were injected throughout each sequence.

### Electrophysiology

Hippocampal brain slices (250 µm thick for whole-cell EPSC and mEPSC recording and 300 µm thick for field and presynaptic mossy fibre bouton recording) were prepared from 21-25 day old ‘wild-type’ (WT; *Slc6a17^+/+^*) or knockout (KO; *Slc6a17^-/-^* mice) following a method described by Bischofberger et al. (2006). Mice were collected on the morning of experiment and promptly euthanised by decapitation without anaesthetic. The brain was rapidly removed and sliced in ice-cold slicing solution then transferred to an incubation chamber containing slicing solution (for the presynaptic recordings) or ACSF (for field and whole-cell EPSC recordings) at 35 – 37°C for up to 30 minutes. Following this time, the incubation chamber was allowed to cool to room temperature. Slices were used for experiments within 6 hours of preparation, or within 3 hours when recording from presynaptic terminals.

Presynaptic mossy fibre boutons were identified in the *stratum lucidum* of CA3 and patch-clamped with a thick-walled borosilicate patch pipette, with a tip resistance of around 8 MΩ, filled with internal solution. Whole-cell mode was obtained and recordings were performed in voltage clamp at -70 mV using an EPC10-double amplifier (HEKA) and Patchmaster software (HEKA, version 2x90.2). Recorded currents were low-pass filtered at 2.9 kHz and digitised at 2 kHz. Recordings were confirmed to be presynaptic by the cells passive membrane properties (high membrane resistance, very fast capacitive currents), active properties (lack of spontaneous synaptic events, firing of a single action potential upon current injection) and post-hoc staining with biocytin included in the patch-pipette (Bischofberger et al., 2006). Glutamine (10 mM) was applied to boutons by 5-second puff application from a pipette placed approximately 50 µm from the terminal, at a repetition rate of one puff per minute, whilst recording the presynaptic currents. Presynaptic glutamine transporter currents were isolated from other currents by a cocktail of pharmacological agents included in bath and puffer solutions (Billups et al., 2013). This comprised: inhibitors of glutamate receptors (40 µM DL-2-amino-5-phospohonopentanoic acid, 10 µM dizocilpine maleate (MK801) and 20 µM NBQX), GABA_A_ receptors (10 µM (-)-bicuculline methochloride), glycine receptors (1 µM strychnine) and Na^+^/K^+^ ion channels (1 µM TTX and 10mM tetraethylammonium chloride). The isolated glutamine transport current amplitudes were recorded in slices from KO mice and WT mice (control) as well as WT slices with 10 mM leucine added to the circulating ACSF.

Evoked EPSCs in CA3 pyramidal cells were obtained by whole-cell voltage-clamp of somas with patch pipettes of resistance of approximately 5 MΩ when filled with internal solution. Following establishment of recordings at -70mV with series resistance was compensated by 70 %, the membrane potential was clamped at +20 mV to record NMDA receptor-mediated EPSCs. Currents were low-pass filtered at 2.9 kHz and digitised at 10 kHz. EPSCs were elicited by constant-current stimulation of presynaptic axons (Digitimer DS2A stimulator) via a concentric bipolar tungsten electrode placed into the mossy fibre pathway. Stimulus strengths ranged from 20 – 230 µA in order to elicit starting EPSC amplitudes of 20 - 60 pA. These EPSCs were confirmed to be the result of mossy fibre stimulation by their sensitivity to group 2/3 metabotropic glutamate receptor stimulation (Figure S8; Kamiya et al., 1996; Lawrence et al., 2004; Jackman et al., 2016). Low-frequency stimulation (LFS; paired pulses, 40 ms apart, repeated every 20 s; Figure 4A) was delivered for 5 minutes, followed by 20 minutes of intermittent high frequency stimulation (HFS; 4 pulses 20 ms apart, repeated every 2 s; Figure 4A). For low frequency-only recordings, LFS was applied continually for 35 minutes.

Spontaneous mEPSCs were recorded by voltage-clamping CA3 pyramidal cells as above, at -70 mV and comprised 1-minute of passive mEPSC recording followed by 5-minutes of HFS (frequencies as above) and another 1-minute passive recording.

In addition to measuring the amplitude of the first EPSC peak or the mEPSCs for comparisons, series and input resistances, paired-pulse ratios and holding currents were measured throughout the recording period. Recordings were excluded if holding current (including compensation) exceeded 600 pA (for evoked EPSCs at +20 mV) or 300 pA (for mEPSCs at -70 mV), or if series resistance changed by more than 35 % or reached 30 MΩ.

Field EPSPs (fEPSPs) were recorded in the *stratum radiatum* of CA1 via a patch pipette of 2-4 MΩ tip resistance, filled with ASCF. Schaffer collaterals were stimulated with low then high frequency as above (Figure 5A). 1 Hz (paired pulses, repeated at 0.5 Hz) was selected as a more appropriate total frequency for HFS at CA1, due to differences in presynaptic surface area and calculated presynaptic depletion rate. Starting fEPSPs were 0.5 to 2 mV in amplitude. Recordings with stimulus artefact amplitude change of more than 5% were excluded.

### Experimental solutions

Slicing solution contained 75 mM sucrose, 87 mM NaCl, 2.5 mM KCl, 25 mM glucose, 1.25 mM NaH_2_PO_4_, 25 mM NaHCO_3_, 7 mM MgCl_2_ and 0.5 mM CaCl_2_ and was continuously bubbled with 95 % O_2_ and 5 % CO_2_.

Experiments were conducted in artificial cerebrospinal fluid (ACSF) heated to 34 ± 0.5°C containing 125 mM NaCl, 2.5 mM KCl, 25 mM glucose, 1.25 mM NaH_2_PO_4_, 25 mM NaHCO_3_, 1 mM MgCl_2_ and 2 mM CaCl_2_ and continuously bubbled with 95 % O_2_ and 5 % CO_2_. Patch pipettes (4.5 - 6 MΩ) were filled with intracellular solution containing: 85 mM CsMeSO_4_, 40 mM HEPES, 10 mM TEA-Cl, 0.008 mM CaCl_2_ and 10 mM EGTA, adjusted to a pH of 7.2 with CsOH.

For evoked EPSC recordings, AMPAR antagonist NBQX (20 µM) and GABA_A_R antagonist bicuculline (10 µM) were added to ACSF to isolate NMDA-mediated currents and were present on the slice for at least 5 minutes prior to the start of recordings. For leucine experiments 10 mM L-leucine was also added. For MeAIB experiment, 20 mM MeAIB was added. Spontaneous mEPSC recordings were conducted in the presence of bicuculline (10 µM) only.

For field recordings, NMDA antagonist APV (40 µM) was added to the ACSF continuously perfusing the slice, and was present for at least 5 minutes prior to the start of recordings in order to prevent changes in synaptic plasticity. For leucine experiments, 1 mM L-leucine was also added. For glutamine ester experiment, glutamine t-butyl ester was added to the incubation chamber to a concentration of 1 mM and slices allowed to incubate for at least 2 hours prior to use. Glutamine ester was also present in the circulating ACSF throughout these recordings.

### Trace fear conditioning

An established method for trace fear conditioning was utilised as outlined in Sellami et al. (2017) and Al Abed et al. (2020).

On the first day, mice were placed in a rectangular Plexiglass conditioning chamber (Imetronic, Pessac, France) which had been wiped down with 70 % ethanol, the lights in the room were on (100 lux) and the stainless-steel rods forming the floor of the chamber was exposed (the ‘conditioning context’). A tone (75 dB, 1 kHz) was played for 30 seconds. This was followed by a ‘trace’ period of 20 or 40 seconds. At the end of this time, a brief (1 s) foot shock (0.2 mA) was delivered by a shock generator (Imetronic, Pessac, France) via the stainless-steel rod flooring of the chamber. This tone-shock pairing was repeated 3 times during a single session. The amount of time spent freezing was measured during the tone and trace periods. On the following day, mice completed two retention tests: a ‘tone test’ and a ‘context test’.

The tone test was similar to one round of the conditioning phase but omitted the foot shock. Rather than setting up the chamber as per the ‘conditioning context’, the sides of the box were obscured with a round bucket insert, wiped with acetic acid for scent and the stainless-steel rod flooring was covered. Lights in the room were dimmed (see Figure 7A). Freezing, defined as the complete lack of movement (excepting respiration) was measured for 2 minutes prior to the tone, while the tone was sounded for the following 2 minutes, during the 20 – 40 second ‘trace’ period after that, and for 20 – 40 seconds following the ‘trace’ period (to make up a total of 2 minutes after the tone).

The context test omitted both the foot shock and the tone but the chamber was set up as per the original ‘conditioning context’ as described above. Freezing was measured during the first 2 minutes, and then between 2-4 minutes and 4-6 minutes.

### Assessment of participation in activities of daily living

Nest building was measured using a method similar to that outlined by Deacon (2006) in which WT and NTT4 KO mice were individually re-housed into freshly prepared cages and provided cotton squares for incorporation into a nest. Mice were left uninterrupted for 24 hours, after which time the nests were visually inspected and a score of 0 – 5 was assigned to reflect the extent of nest assembly with 0 indicating that the cotton remained completely intact and 5 indicating the construction of a complete nest with a cup-shaped base and sides.

### Assessment of locomotion and anxiety

Locomotion and anxiety can both be assessed using an open field test, whereby mice are placed into a high-walled open arena and their movements within the space recorded over a number of minutes. To measure distance travelled and thigmotaxis during a 4-minute period, WT and NTT4 KO mice were placed, one at a time, into the centre of a white plastic arena 38.5 cm in diameter with 40cm high walls. The arena was brightly lit at 100 lux via an overhead desk lamp and a webcam was also mounted above the arena in order to record the movements of the mouse within the space.

### Comparison of social preference

To assess social interaction in mice, a two-chamber social preference test was used, based on the protocol described by Kaidanovich-Beilin et al. (2011). WT and NTT4 mice were individually placed into the centre of a dual chambered enclosure in which one side contained a novel object (a LEGO figure) and the other contained a sex-matched mouse with whom the subject mouse had not previously interacted. The novel mouse was contained within a ventilated secondary container. A suspended webcam was used to record the subject mouse’s movements in the chamber, and the duration of interactions with both the novel object and novel mouse were later measured from these recordings.

### Data analysis and statistics

Electrophysiology data was recorded with Patchmaster software (HEKA. version 2x90.2). For evoked EPSCs and fEPSPs, data extraction and averaging were performed using MATLAB (Version 9.3) and for spontaneous mESPCs, AxoGraph X (Version 1.8.0) was used to detect mEPSCs using a pre-defined template to calculate amplitude and frequency averages.

For behavioural tests, time spent freezing was measured during the relevant time points and recorded and averaged in StatView (Version 5.0.1) spreadsheets.

^13^C-NMR and LC-MS, and synaptosome amino acid uptake data were plotted into and averaged using Microsoft Excel spreadsheets.

In the open field test, movements were tracked and analysed automatically using a MATLAB-based detection program (Zhang et al., 2020) adapted for use in a circular chamber (Parkinson et al., 2024). Total distance travelled was compared between groups, as was thigmotaxis; measured as the percentage of time spent in the outer 2.5 inches (6.35cm) of the arena.

Social preference was calculated as in (Al Abed et al., 2024), as the ratio of time spent interacting with the novel mouse, over the total interaction time; comprising time spent interacting with both the other mouse and the novel object (mouse / mouse + object).

For fear conditioning tone test data, a freezing index was calculated by dividing the time spent freezing during the tone and trace, minus freezing before the tone (%) by the time spent freezing during tone, trace and pre-tone combined (%).

Data were then analysed in RStudio (Version 4.0.2) using the linear mixed effects model package ‘nlme’ (Pinheiro and Bates, 2000; Pinheiro et al., 2020) with cell or animal as a random factor. mEPSC sample distribution analysis was undertaken using an asymptotic two-sample Kolmogorov-Smirnov test, also in RStudio. All data are presented as mean ± SEM. Statistical significance is indicated by p < 0.05.

## Acknowledgments

We thank the members of the Billups lab for discussion and suggestions, members of the Australian Phenomics Facility for animal care and husbandry, and the Mark Wainwright Analytical Centre, UNSW for technical support. We would also like to thank Mrs Jing Gao, Dr Nay-Chi Khin and Dr Jenna Lowe for their assistance in the generation of the knockout mice. This work was supported by grants from the National Health and Medical Research Council, Australia (grant GNT1105857 to B.B and S. Bröer), Australian Research Council (grant DP180101702 to S. Bröer, C.D.R and B.B), the National Collaborative Infrastructure Strategy via Phenomics Australia (to G.B.) and internal funding from the John Curtin School of Medical Research.

## Author contributions

A.L.N., A.S.A., S.R.H., G.B., C.D.R., S. Bröer and B.B. designed research; A.L.N., A.S.A., S.R.H., A.D., G.G., A.B., G.B., C.D.R., S. Bröer and B.B. performed research; A.B., S. Balkrishna, G.B., N.D. and B.B. contributed new research and analytical tools; A.L.N., A.S.A., S.R.H., A.D., G.G., A.B., C.D.R., S. Bröer and B.B. analyzed data; A.L.N. and B.B. wrote the paper. All authors approved of the final version of the manuscript.

## Data Availability

Data associated with this study can be found at Mendeley Data doi: 10.17632/rfphzt3wjm.1

