## Supporting Information for "Glutamine Transport via Neurotransmitter Transporter 4 (NTT4, SLC6A17) Maintains Presynaptic Glutamate Supply at Excitatory Synapses in the Central Nervous System"

**This PDF file includes:**
Figures S1 to S8

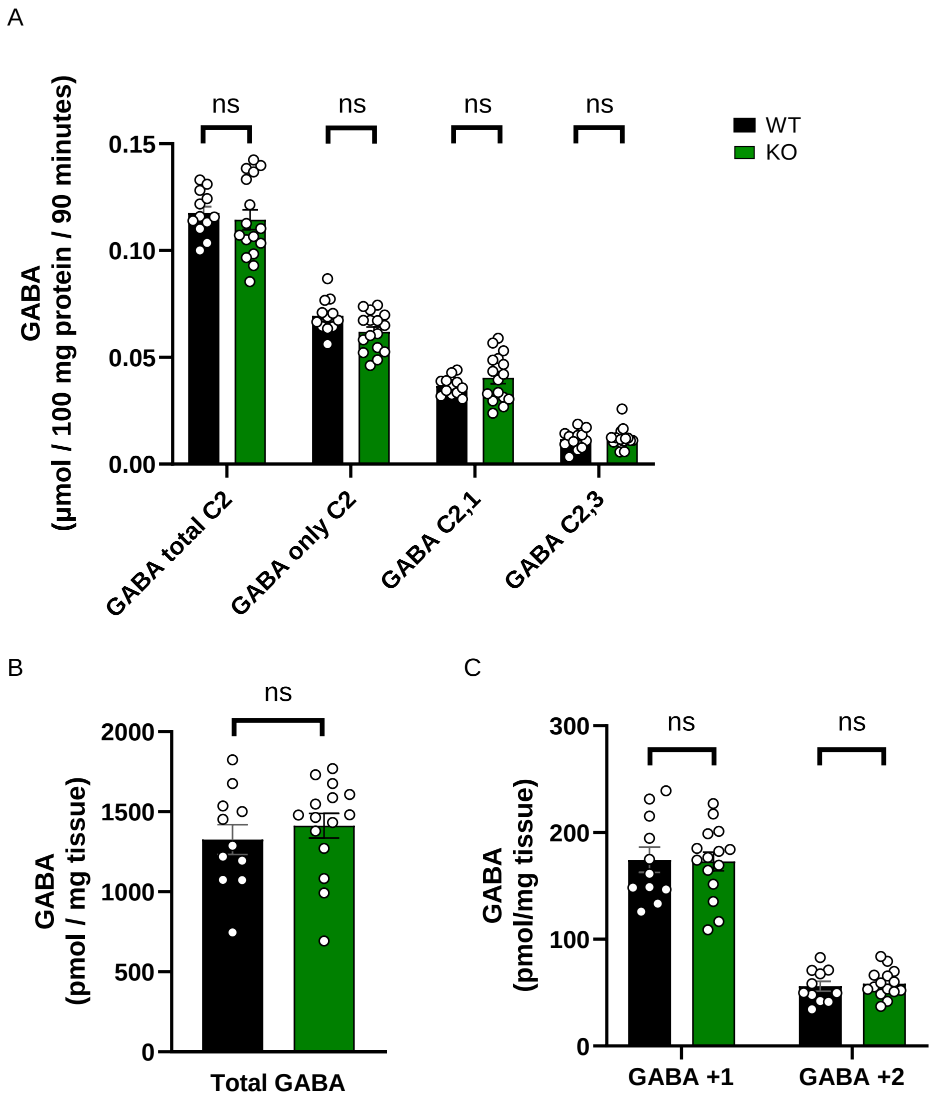


**Figure S1. GABA production is not reduced in NTT4 KO mice.**

**(A)** In cortical samples analysed with NMR spectroscopy, isotopically enriched GABA in NTT4 KO samples (green) is not different to control (WT, black). This includes all GABA labelled at the second carbon (total C2), GABA derived from neuronal metabolism of glucose, singly labelled at C2 (only C2), and GABA derived from astrocytic metabolism of acetate, labelled at both the first and second carbons (C2,1) and second and third carbons (C2,3) **(B)** Similarly, in hippocampal samples analysed with LCMS, there is no difference in total GABA between KO (green) and control (WT; black) **(C)** In hippocampal samples analysed with LCMS, there is no difference in singly-labelled GABA (GABA +1), resulting primarily from metabolism of glucose in neurons, in KO (green) vs control (WT; black), nor of doubly-labelled GABA (GABA +2) resulting primarily by metabolism of acetate in astrocytes.

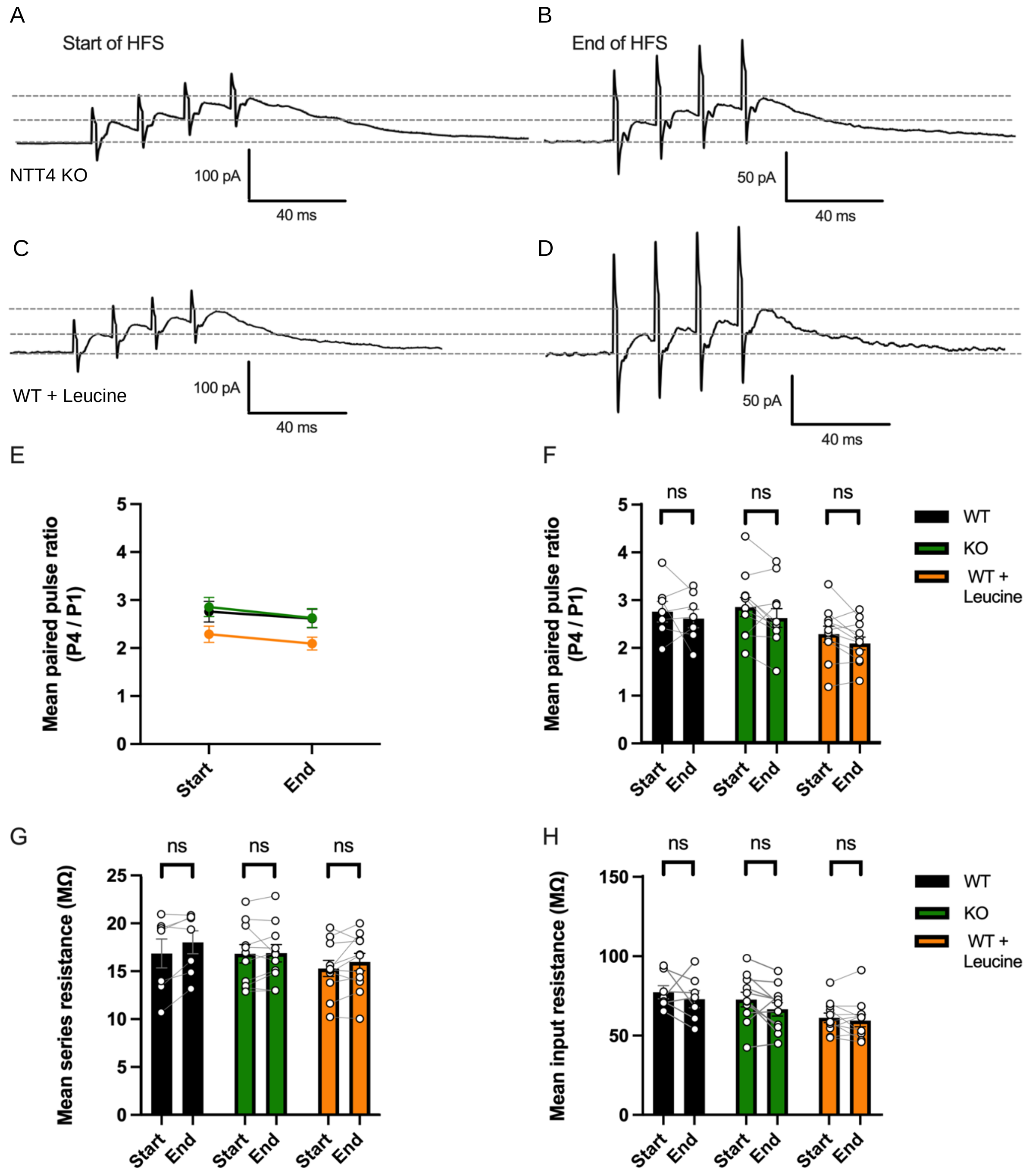


**Figure S2. EPSC amplitude reduction in NTT4 KO cells and WT cells treated with leucine is not accompanied by changes in paired-pulse ratio, nor series or input resistance.**

**(A)** Example raw data traces from one NTT4 KO cell recording at the beginning, and **(B)** end of high-frequency stimulation (HFS), with the latter re-scaled to highlight the unchanged paired-pulse ratio. Grey dotted lines highlight the first and last EPSC amplitudes for comparison. **(C)** Example raw data traces from one WT + leucine recording at the beginning, and **(D)** end of HFS, with the latter re-scaled to highlight the unchanged paired-pulse ratio. Grey dotted lines highlight first and last EPSC amplitudes for comparison. **(E)** Mean paired-pulse ratios for WT (black), NTT4 KO (green) and WT + leucine (orange) recordings at the start and end of HFS showing no difference between groups. **(F)**

Mean paired-pulse ratios for WT (black), NTT4 KO (green) and WT + leucine (orange) recordings show no difference between start and end of HFS in any group **(G)** Mean series resistance for WT (black) and NTT4 KO (green) and WT + leucine (orange) recordings at start and end of recording shows no change in series resistance in any group. **(H)** Mean input resistance for WT (black), NTT4 KO (green) and WT + leucine (orange) groups at start and end of recording, shows no change from start to end on either group. Error bars show SEM and data points indicate individual recordings. ‘ns’ denotes no significant difference ($p > 0.05$).

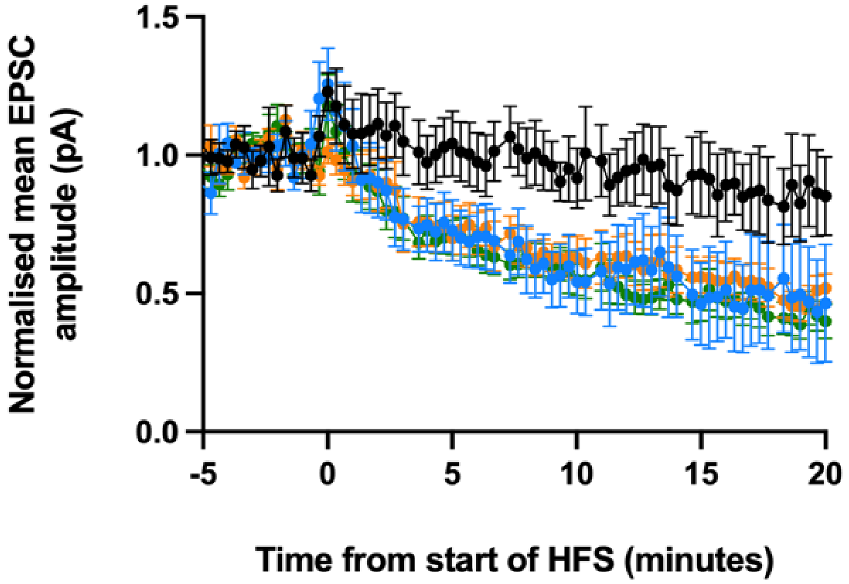


**Figure S3. The effect of leucine is via its action on NTT4.**

EPSC amplitudes (normalised to the 5 minutes of LFS for each group) can be seen to reduce equally during HFS in cells from KO animals with leucine present (blue), in WT animals with leucine present (orange) and in cells from NTT4 KO animals (green) without leucine. The lack of additional effect on leucine in NTT4 KO mice indicates that the inhibition of EPSCs by leucine is via its effect on NTT4. There is no reduction in amplitude in cells from WT (control) animals (black). Data points show mean EPSC amplitudes for each group, in LFS these are for each sweep and for HFS is the average of 10 sweeps. Gaps are present after each 100 sweeps during which time series resistance compensation is switched off in order to gauge the uncompensated series resistance and input resistance, but stimulation frequency is maintained throughout. Error bars indicate SEM.

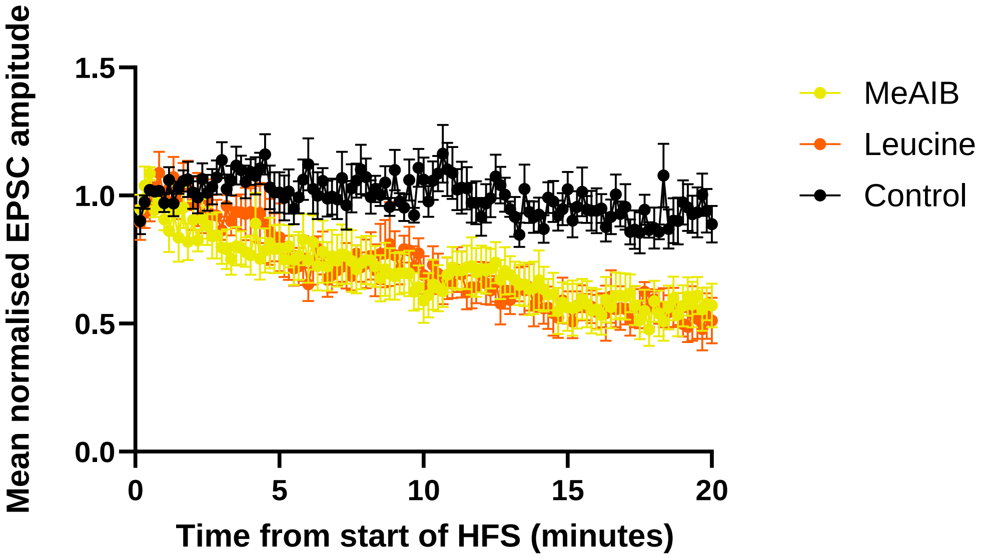


**Figure S4. System A (SNAT1/2) transporters do not provide an appreciable supply of glutamine to presynaptic terminals during HFS.**

(A) EPSC amplitudes (normalised to the 5 minutes of HFS for each group) can be seen to reduce equally in cells from WT animals with leucine present (orange) and in cells from WT animals that have been treated with the amino acid analogue MeAIB (yellow), indicating a lack of significant role for SNAT1/2 in replenishing presynaptic glutamate at this synapse There is no reduction in amplitude in cells from WT (control) animals (black). Data points show mean EPSC amplitudes for each group, averaged over 10 sweeps. Error bars indicate SEM.

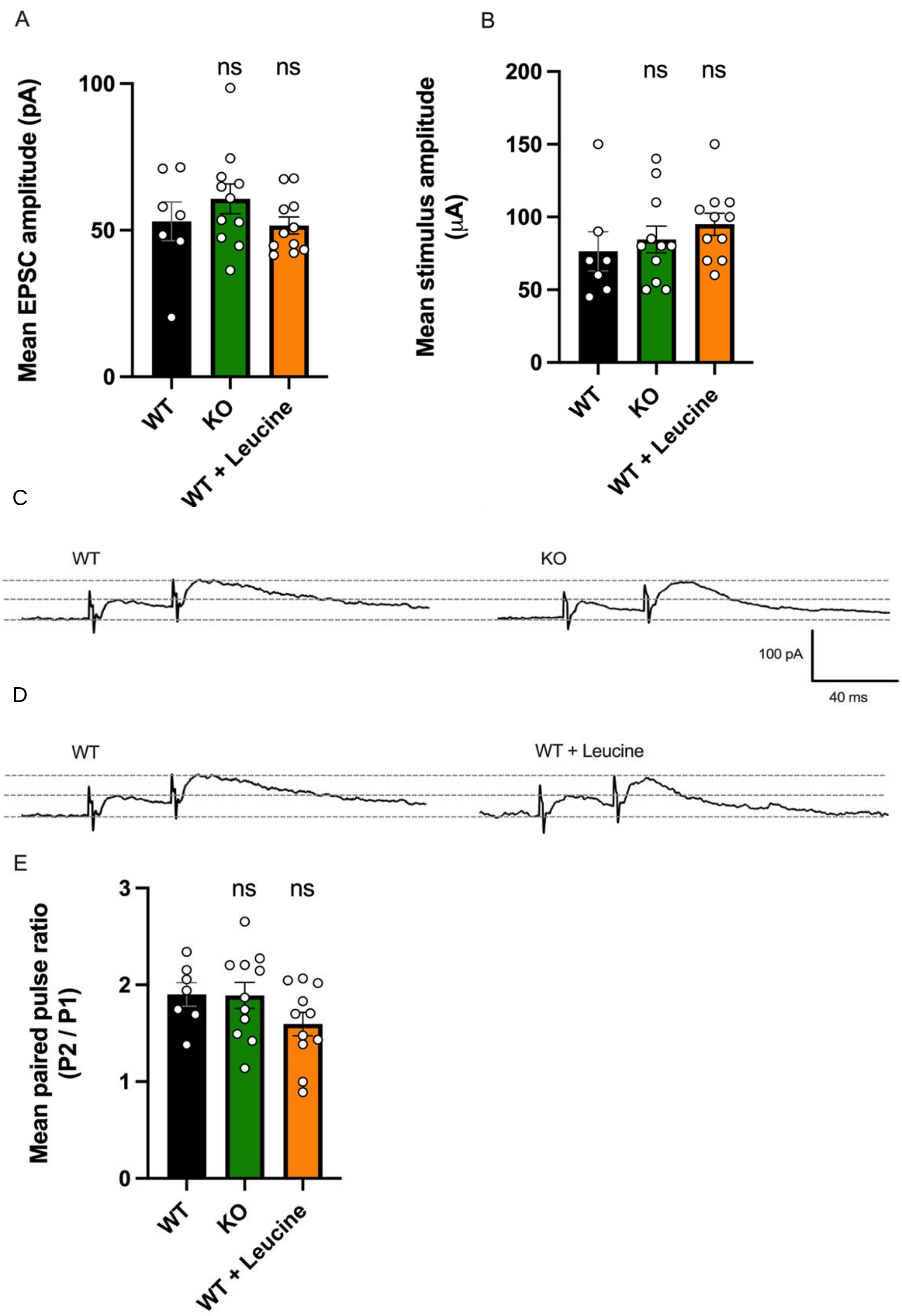


**Figure S5. EPSCs evoked in NTT4 KO cells and WT cells with leucine were no different to WT control cells during low-frequency stimulation (LFS).**

**(A)** Mean EPSC amplitudes during LFS were similar between WT control (black), NTT4 KO (green) and WT cells in the presence of 10 mM leucine (orange). **(B)** Amplitude of the stimulus current pulses used to elicit the EPSCs did not differ between WT control (black), KO (green) and WT + leucine (orange). **(C)** Raw data traces from whole-cell recordings in a WT cell (left) and an NTT4 KO cell (right), showing evoked EPSCs during LFS. Baseline and amplitude of the first and second peaks are highlighted with dashed grey lines for comparison. **(D)** Same as for (C) but showing raw data traces from a WT cell (left) and a WT cell in the presence of 10 mM leucine (right). **(E)** Paired-pulse ratios in NTT4 KO and WT cells in the presence of leucine were not different to WT control during LFS. Error bars indicate SEM and data points show individual recordings. 'ns' indicates no significant difference ($p > 0.05$).

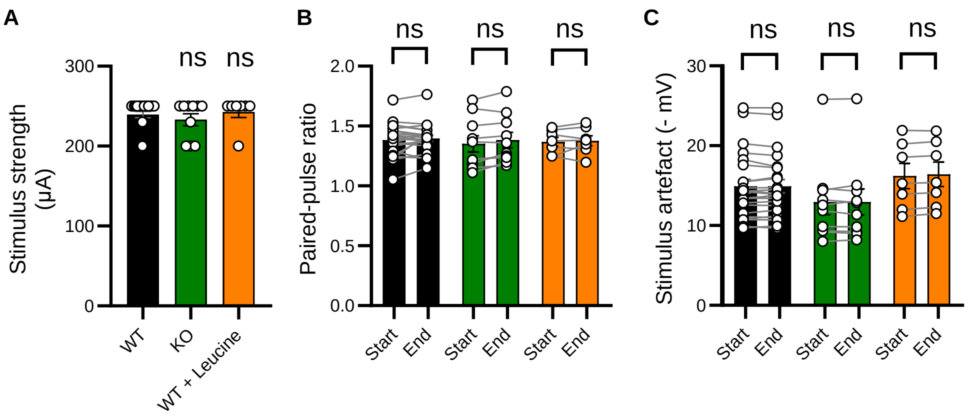


**Figure S6. Reduction of fEPSP amplitude in KO and leucine-treated WT slices during HFS is not accompanied by differences in stimulation strength, paired-pulse ratio change or stimulus artefact amplitude.**

**(A)** Stimulus strengths used to elicit fEPSPs in NTT4 KO slices (green) and WT slices treated with leucine (orange) did not differ from the strengths used for WT control recordings (black). **(B)** Paired-pulse ratio did not change in recordings from NTT4 KO (green) or WT + leucine slices (orange) and were comparable to those in WT control recordings (black) **(C)** Stimulus artefacts did not change over time in WT (black), NTT4 KO (green), or WT + leucine recordings (orange), showing effective stimulation was unchanged throughout the experiment. Error bars show SEM and data points show individual recordings. ‘ns’ denotes no significant difference ($p > 0.05$).

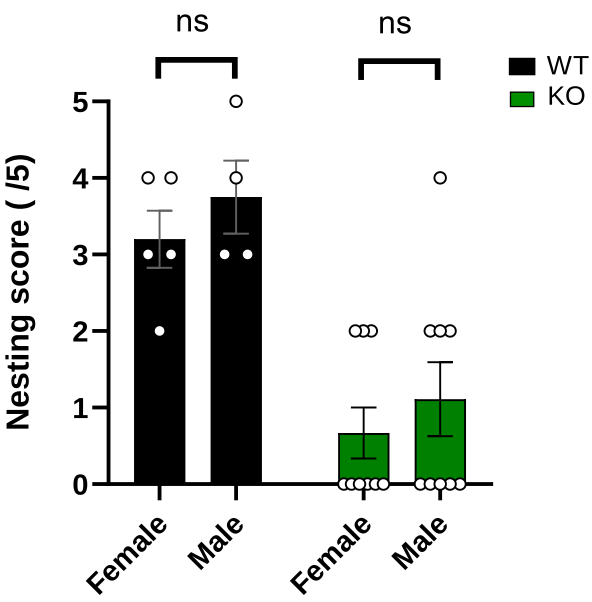


**Figure S7. Nesting scores in NTT4 and WT mice did not differ by sex**

Nesting scores on a scale of 0 – 5 divided by sex. Scores from male vs female mice did not differ in either WT control (black) or NTT4 KO (green) groups. Error bars show SEM and data points show individual recordings. ‘ns’ denotes no significant difference ($p > 0.05$).

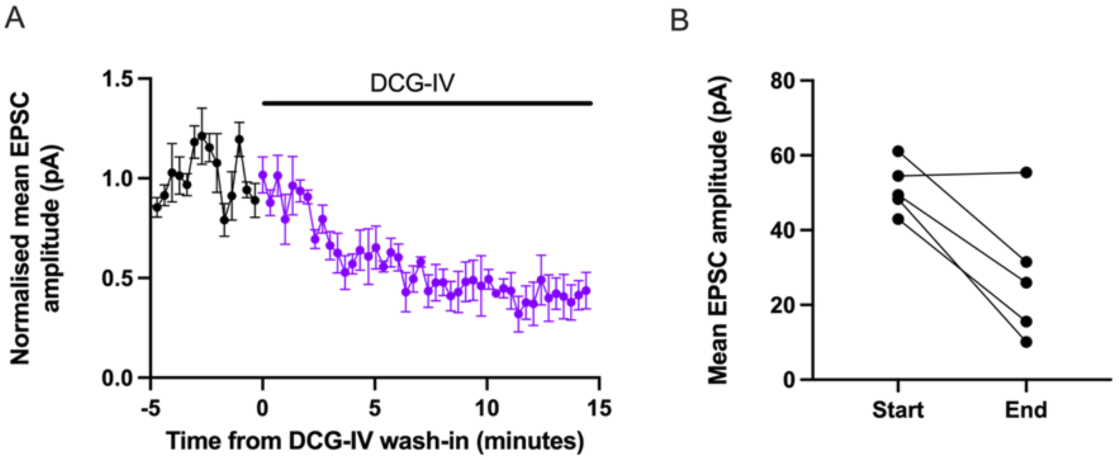


**Figure S8. Inhibition of CA3 EPSCs by mGluR activation confirms mossy fibre stimulation.**

**(A)** Normalised mean NMDA-mediated EPSC amplitudes recorded at +20 mV from WT CA3 cells receiving DCG-IV-sensitive presynaptic inputs (n = 4), before (black) and after the addition of mGluR$_{2/3}$ agonist DCG-IV (1 µM, purple). A reduction of around half of the original amplitude is evident after 10 – 15 minutes post wash-in. **(B)** Addition of DCG-IV led to a reduction in EPSC amplitude in 4 out of 5 cells measured before and after addition of the drug to the circulating ACSF. Data points show the mean EPSC amplitude for each cell before and after addition of DCG-IV. The greater than 50% reduction in EPSC amplitude in the majority of recordings is consistent with mossy fibre stimulation (Lawrence et al., 2004).
